# Improvement of Behavioral Deficits in a Mouse Model of Fragile X Syndrome by Restoring α7 nAChR Hypofunction Associated with Aberrant Ly6H Expression

**DOI:** 10.1101/2024.11.01.621616

**Authors:** Sarah Goebel, Dylann Cordova-Martinez, Vytas K. Verselis, Anna Francesconi

## Abstract

Fragile X Syndrome (FXS) is the most common form of inherited intellectual disability and often accompanied with debilitating pathologies including seizures and hyperactivity. FXS arises from a trinucleotide repeat expansion in the 5’ UTR of the *FMR1* gene that silences expression of the RNA-binding protein FMRP. Despite progress in understanding FMRP functions, the identification of effective therapeutic targets has lagged and at present there are no viable treatment options. Here we identify the α7 nicotinic acetylcholine receptor (nAChR) as candidate target for intervention in FXS. In the early postnatal hippocampus of *Fmr1* knockout (*Fmr1*^KO^) mice, an established pre-clinical model of FXS, the α7 nAChR accessory protein Ly6H is abnormally distributed, showing enrichment at the neuronal surface and mislocalization in dendrites. Ly6H, a GPI-anchored protein, binds α7 nAChRs with high affinity and can limit α7 nAChR surface expression and signaling. We find that α7 nAChR-evoked Ca^2+^ responses are dampened in immature glutamatergic and GABAergic *Fmr1*^KO^ neurons compared to wild type. Knockdown of endogenous Ly6H in *Fmr1*^KO^ neurons is sufficient to rescue dampened α7 nAChR Ca^2+^ responses *in vitro*, providing evidence of a cell-autonomous role for Ly6H aberrant expression in α7 nAChR hypofunction. In line with intrinsic deficits in α7 nAChR activity in *Fmr1*^KO^ neurons, *in vivo* administration of the α7 nAChR-selective positive allosteric modulator PNU-120596 improved spatial memory and reduced hyperactivity and seizure severity in adolescent *Fmr1*^KO^ mice. Taken together, our *in vitro* mechanistic findings and *in vivo* rescue studies implicate α7 nAChR hypofunction in FXS pathology.

## INTRODUCTION

Fragile X Syndrome (FXS) is the most prevalent form of inherited intellectual disability with approximately 1/7,000 males and 1/11,000 females diagnosed (*1*). FXS is often co-morbid with associated pathologies including seizures in childhood (∼ 18% of males, 7% females), attention deficit hyperactivity (∼ 80%) and autism (∼ 30%) (*2, 3*). FXS arises from the abnormal expansion of a trinucleotide repeat in the 5’ UTR of the X-linked *FMR1* gene resulting in its silencing by hypermethylation and global loss of expression of the encoded Fragile X messenger ribonucleoprotein (FMRP) (*4, 5*). FMRP is an RNA-binding protein that regulates translation, trafficking, and stability of ∼ 8% of brain-expressed RNAs (*5*). Studies utilizing the *Fmr1* knockout (*Fmr1*^KO^) mouse have revealed that loss of FMRP is associated with dendritic spine dysgenesis, deficits in forms of synaptic plasticity, and hyperexcitability (*6*). Despite remarkable progress in understanding the cellular and synaptic dysfunctions arising from *FMR1* silencing, there are currently no treatment options for patients beyond palliative measures, underscoring the urgent need to identify avenues for therapeutic intervention.

Acetylcholine (ACh) is a key neuromodulator/neurotransmitter that through widespread cholinergic fiber innervation plays an important role in regulating critical periods of brain development driven by sensory and experience-dependent activity (*7–9*). ACh actions are mediated in part by nicotinic ACh receptors (nAChRs), a family of ligand-gated ion channels broadly expressed in the central nervous system (CNS). Amongst nAChRs, the homopentameric α7 nAChRs play a crucial role during early postnatal development in the maturation of excitatory and inhibitory neurons, the refinement of their synaptic connections, and the establishment of functional neural circuits (*9–13*). Although alterations in signaling by G protein-coupled muscarinic ACh receptors were reported in *Fmr1*^KO^ mice (*14, 15*), whether α7 nAChR activity is compromised in FXS remains untested despite evidence of its role in many of the processes that go awry in individuals with FXS including seizures, hyperactivity, and hypersensitivity to sensory stimuli (*16–19*). The critical impact of α7 nAChRs on neuronal function and maturation is attributable, in large part, to its high Ca^2+^ permeability (*20*) that can initiate downstream signaling and affect neurotransmitter release. Tight control of α7 nAChR activity is essential to limit excessive cytotoxic Ca^2+^ influx and is achieved in part through regulated biogenesis and surface export (*21, 22*). These processes are regulated by accessory proteins, including chaperones and auxiliary proteins such as Ly6/uPAR proteins. The glycosylphosphatidylinositol (GPI)-anchored Ly6/uPAR proteins are an integral component of the nAChR signaling unit (*23*). They bind to nAChRs *via* a three-finger domain structurally related to that of α-Bungarotoxin which binds nAChRs with high affinity (*23*). Seminal studies established a role of members of the Ly6/uPAR protein family as endogenous modulators of nAChR activity in the CNS by enhancing receptor desensitization and reducing surface expression (*24–27*). We previously reported a genome wide proteomic screen of the adult *Fmr1*^KO^ mouse brain that found Lymphocyte Antigen 6 Family Member H (Ly6H), a member of the Ly6/uPAR family, to be selectively depleted from membrane fractions containing lipid rafts (*28*). Ly6H, a brain enriched protein expressed in neurons, is of particular interest as it binds to and regulates α7 nAChR function by restraining its activity through direct interaction at the plasma membrane (*29*) and by limiting its surface expression (*26, 30*).

In this study, we report that immature hippocampal *Fmr1*^KO^ neurons exhibit aberrantly elevated Ly6H surface expression, particularly in the soma and proximal dendritic regions. Concomitant with altered Ly6H expression and congruent with its functions, we find reduced evoked Ca^2+^ responses of α7 nAChRs in *Fmr1*^KO^ hippocampal neurons of both glutamatergic and GABAergic lineage that can be corrected by downregulation of Ly6H. We further demonstrate that early pharmacological enhancement of α7 nAChR activity in adolescent *Fmr1*^KO^ mice improves hippocampal-based object location memory and reduces hyperactivity and the severity of audiogenic seizures. Thus, our results identify α7 nAChR hypofunction as a critical deficit arising from loss of FMRP and provide support for the therapeutic benefits of augmenting α7 nAChR activity in FXS.

## RESULTS

### Increased surface expression and altered localization of Ly6H in *Fmr1*^KO^ hippocampal neurons

In previous work, we used isobaric tags (iTRAQ) and mass spectrometry applied to adult wild type (WT) and *Fmr1*^KO^ forebrain to characterize the protein composition of detergent resistant membranes (DRMs) that contain lipid rafts (*28*). Unlike most other GPI-anchored proteins identified in the screen, Ly6H (**Figure 1A**) was found to be depleted in DRMs from cortex and hippocampus of *Fmr1*^KO^ mice relative to WT without substantial changes in total Ly6H protein levels. GPI-anchored proteins are normally transported to and retained in lipid rafts, and the altered membrane distribution of Ly6H suggested potential defects in its trafficking and or cellular localization. Given that postnatal days 7-8 (P7-8) is the age at which Ly6H expression becomes relatively abundant in the hippocampus (**Figure S1A-B**), we examined Ly6H expression at the cell surface in hippocampal tissue slices obtained from WT and *Fmr1*^KO^ pups at P7-8 (**Figure 1B**). The hippocampal slices were live labeled with biotin and biotinylated proteins were precipitated with streptavidin and analyzed by Western blot. The proportion of surface expressed Ly6H proved to be upregulated in the *Fmr1*^KO^ hippocampus, with mean 2.3-fold increase relative to WT, despite total Ly6H abundance remaining comparable between genotypes (**Figure 1B-C**). The transcriptional RNA-sequencing (RNA-seq) profiles of single cells in the murine brain indicate that Ly6H mRNA is predominantly expressed in neurons (*31*), including principal neurons in all subregions of the hippocampus (*32*). To visualize Ly6H protein expression, Ly6H was immunolabeled in hippocampal slices obtained from WT and *Fmr1*^KO^ littermates at P7 and P22. At both ages, labeled Ly6H appeared punctate in both the pyramidal cell layer, marked by nuclear labeling with DAPI, and the neuropil (**Figure 1D**). The density of Ly6H puncta was higher in the *Fmr1*^KO^ compared to WT both in the CA1 pyramidal layer (**Figure 1E**) and in the proximal region of the *stratum radiatum* (extending 40 μm from the pyramidal layer) that harbors the apical dendrites of pyramidal neurons (**Figure 1F**). To examine Ly6H dendritic localization more closely, we used *Fmr1*^KO^ and WT primary hippocampal neurons which develop a complex dendritic arbor by 7 days *in vitro* (d.i.v.). This strategy allowed application of Ly6H antibody to non-permeabilized cells to visualize surface Ly6H, followed by post-permeabilization immunolabeling of the dendritic marker Microtubule-Associated Protein 2 (MAP2). The soma and the dendritic arbor were studded with Ly6H puncta in both genotypes (**Figure 1G-H**) but the area occupied by labeled Ly6H was ∼ 1.5-fold greater in the *Fmr1*^KO^ soma compared to WT (**Figure 1I**). Notably, Ly6H-positive puncta appeared unevenly distributed in *Fmr1*^KO^ dendrites and more densely clustered within 10 μm from the soma compared to WT (**Figure 1H** and **1J**). As observed in hippocampal slices, despite altered surface expression, total Ly6H protein abundance in primary hippocampal neurons was comparable between genotypes (**Figure S2A-B**). Together, these findings indicate that Ly6H, an accessory protein to α7 nAChR, is abnormally localized and exhibits increased surface clustering in the proximal dendritic regions and soma of immature *Fmr1*^KO^ hippocampal neurons.

**Figure 1.**
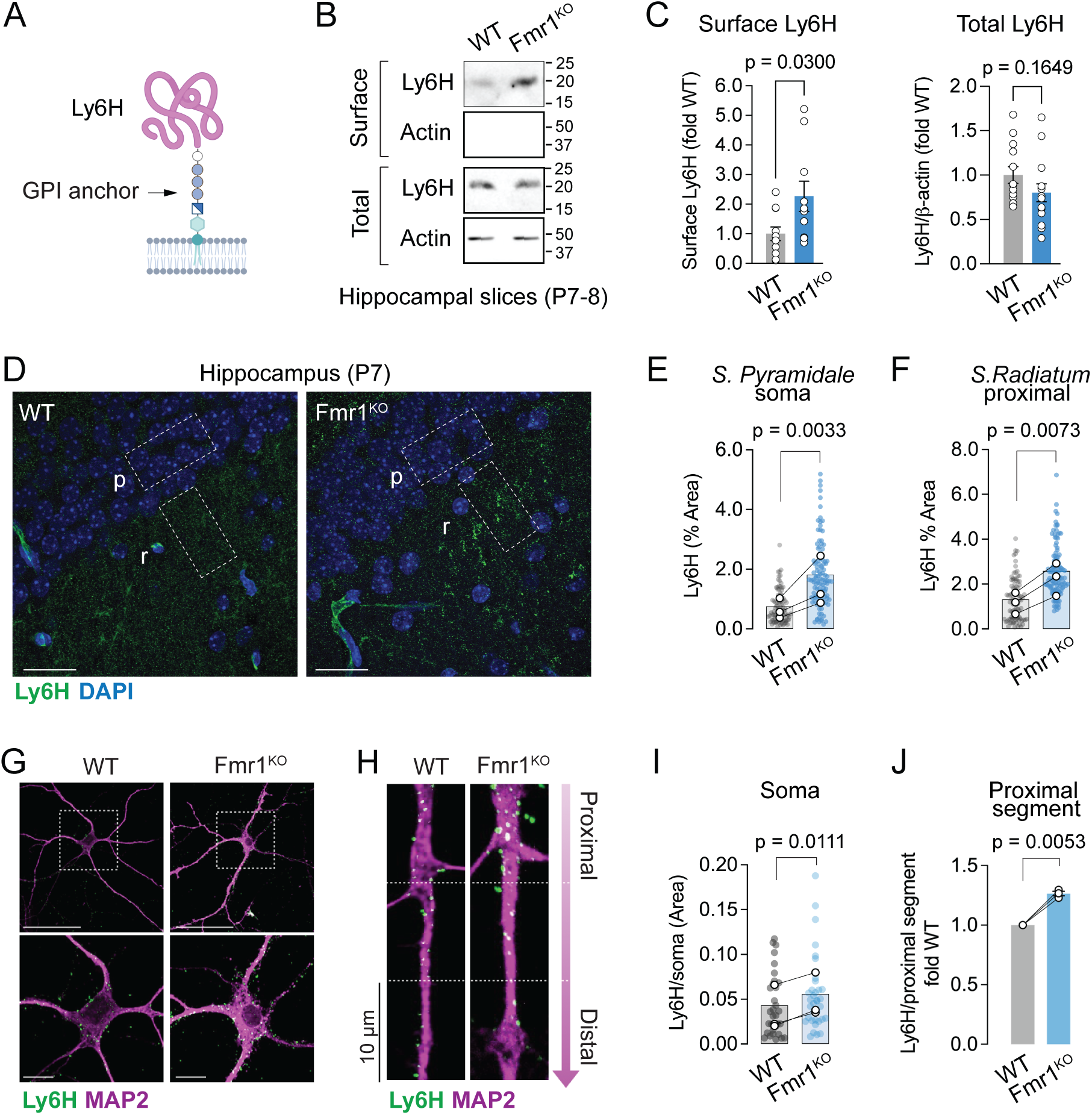
Increased surface expression and altered localization of Ly6H in *Fmr1*^KO^ hippocampal neurons. **(A)** Schematic of Ly6H structure (created with BioRender). (**B**) Representative Western blots of surface (biotinylated) and total Ly6H in hippocampal slices (Actin, loading control). (**C**) Quantification of normalized surface (left, WT n = 11, *Fmr1*^KO^ n = 10 mice) and total Ly6H (right, WT n = 13, *Fmr1*^KO^ n = 14 mice); unpaired t-test. (**D**) Representative confocal images of Ly6H and nuclei stained with DAPI in hippocampal slices. Shown is area CA1: *p*, pyramidal layer, *r*, *stratum radiatum*. Outlined regions (20 x 40 µm) are representative of those used for quantification; scale bars, 25 µm. (**E**) Quantification of Ly6H in the CA1 pyramidal layer containing soma; percentage of Ly6H area in regions randomly selected in P7 and P22 mice (N = 3 mice per genotype). Filled symbols correspond to all regions analyzed (WT n = 91; *Fmr1*^KO^ n = 110) and outlined symbols are the means of matched littermates. Mean ± SEM: WT 0.66 ± 0.19, *Fmr1*^KO^ 1.5 ± 0.48, N=3, ratio paired t-test. (**F**) Quantification of Ly6H in the *stratum radiatum*; percentage of Ly6H area in regions randomly selected as in (D). Filled symbols are all regions analyzed (WT n = 94; *Fmr1*^KO^ n = 113) and outlined symbols are the means of matched littermates. Mean ± SEM: WT 1.2 ± 0.27, *Fmr1*^KO^ 2.2 ± 0.42, N=3, ratio paired t-test. (**G**) Representative confocal images of surface Ly6H in the somatodendritic region of primary hippocampal neurons at d.i.v. 7 with boxed regions shown magnified below; scale bars 50 and 10 µm, respectively. (**H**) Representative confocal images of surface Ly6H in proximal regions of primary dendrites of d.i.v. 7 hippocampal neurons. The first three proximal segments (closest to the soma) are shown, each 10 µm in length. (**I**) Quantification of somatic Ly6H puncta area normalized to soma area from images as in (G). Filled symbols are all soma analyzed (WT n = 33, *Fmr1*^KO^ n = 36), outlined symbols are the means from individual matched cultures (N = 3). Mean ± SEM: WT 0.036 ± 0.015, *Fmr1*^KO^ 0.051 ± 0.014, paired t-test. (**J**) Quantification of fold difference of normalized Ly6H puncta area in the first proximal segment from images as in (H). KO/WT ratio, mean ± SEM, 1.3 ± 0.019 from N = 3 cultures in which n = 137 and n = 126 segments were quantified for WT and *Fmr1*^KO^, respectively; one sample t-test.

### Evoked α7 nAChR calcium responses are dampened in *Fmr1*^KO^ neurons

Ly6H was shown to regulate α7 nAChR function by restraining its activity at the plasma membrane through direct association (*29*) and also by limiting surface expression of mature α7 nAChRs (*26, 30*). These findings prompted us to consider the possibility that the aberrant localization of Ly6H we observed in *Fmr1*^KO^ neurons may lead to intrinsically altered α7 nAChR activity. The α7 nAChR is one of the most abundant nAChRs in the brain and is predominant in the hippocampus, where its expression peaks in the first two postnatal weeks in rodents (*33, 34*). α7 nAChRs are expressed in excitatory neurons and are particularly enriched in GABAergic neurons, including during the early immature stage when GABA is excitatory (*34–36*). To examine α7 nAChR responses, we sought to take advantage of α7 nAChR’s very high Ca^2+^ permeability that can produce rapid increases in cytosolic Ca^2+^ upon receptor activation. We first measured α7 nAChR responses in WT primary hippocampal neurons, loaded with the ratiometric Ca^2+^ indicator Fura-2AM, that were evoked by bath application of the α7 nAChR-selective agonist PNU-282987 (100 nM) (*37*) together with the α7 nAChR-selective positive allosteric modulator (PAM) PNU-120596 (3 µM) (*16*). The PAM PNU-120596 is commonly used to facilitate Ca^2+^ detection by preventing the very rapid desensitization of α7 nAChRs (*20*). Stimulation of α7 nAChRs with a combination of agonist and PAM produced a robust and rapid rise in intracellular Ca^2+^ (**Figure 2A-B**) that could be blocked by the selective antagonist methyllycaconitine (10 nM), confirming that responses were mediated by α7 nAChRs (**Figure 2C**). Since α7 nAChR activity and expression in the hippocampus were shown to be higher in GABAergic cells compared to glutamatergic neurons (*35, 38*), we used post-hoc immunolabeling of glutamic acid decarboxylase (GAD65) to allow assignment of the evoked Ca^2+^ transients to either GABAergic or glutamatergic neurons (**Figure 2D-E**). In WT neurons, PNU-282987 near its EC_50_ (100 nM) (*39*), but not at 10-fold lower concentration (10 nM), consistently produced evoked Ca^2+^ responses in both glutamatergic and GABAergic cells (**Figure S3A-D**). GAD65-positive (GAD65+) GABAergic cells exhibited significantly stronger Ca^2+^ responses compared to glutamatergic cells negative for GAD65 (GAD65-), both in regard of higher peak amplitudes (**Figure 2F-G**) and overall integrated areas of the Ca^2+^ responses relative to baseline (**Figure 2H-I**). This result is congruent with higher α7 nAChR expression reported in GABAergic neurons.

**Figure 2.**
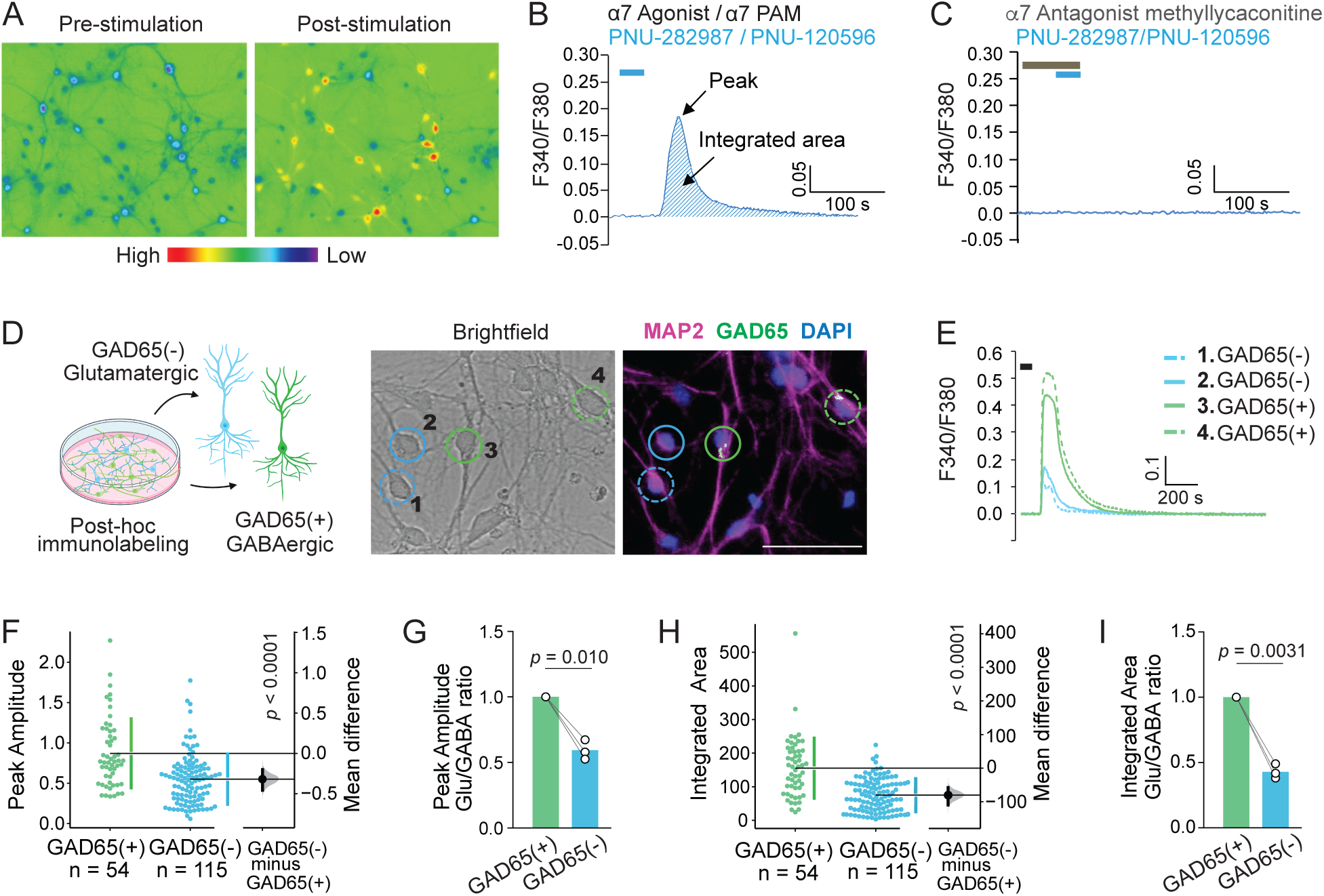
Immature WT hippocampal GABAergic neurons show stronger α7 nAChR-mediated Ca^2+^ responses compared to glutamatergic neurons. (**A**) Representative images of Ca^2+^ rise in WT hippocampal neurons before (pre) and after (post) bath-applied α7 nAChR agonist together with PAM (as in B); high/low heatmap visualizes Ca^2+^ level. (**B)** Representative trace of a Ca^2+^ transient evoked by PNU-282987 (100 nM) with PNU-120596 (3 μM) in a WT hippocampal neuron (d.i.v. 9); bar indicates time and duration of drug application (**C**) The Ca^2+^ rise induced by PNU-282987 with PNU-120596 is blocked by methyllycaconitine; bars indicate time and duration of drug application. (**D**) Schematic of the strategy to distinguish GABAergic (GAD65+) *vs*. glutamatergic (GAD65-) cells (created with BioRender), and representative images of d.i.v. 7 GAD65(+) (green circles) and GAD65(−) (blue circles) hippocampal neurons; scale bar 50 µm. (**E**) Responses evoked by PNU-282987 with PNU-120596 in cells circled in (D). (**F**) Quantification of peak amplitude of α7 nAChR responses in GAD65(+) and GAD65(−) neurons from N = 3 experiments. Gardner-Altman estimation plot; group values (left axes) and mean difference (right floating axes) shown as a bootstrap sampling distribution. The mean difference is indicated by the dot and the 95% confidence interval is indicated by the error bar; two-sided permutation t-test. (**G**) Ratio of mean peak amplitudes; GAD65(−)/GAD65(+) mean ± SEM, 0.59 ± 0.043, N = 3, one sample t-test. (**H**) Quantification of the integrated area in GAD65(+) and GAD65(-) neurons from N = 3 experiments; Gardner-Altman estimation plot, two-sided permutation t-test. (**I**) Ratio of mean integrated area; GAD65(-) /GAD65(+) mean ± SEM, 0.43 ± 0.032, N = 3, one sample t-test.

Having established the properties of evoked α7 nAChR responses in these two neuronal types, we next sought to characterize α7 nAChR function in *Fmr1*^KO^ hippocampal neurons over the course of a critical maturation time window *in vitro* from d.i.v. 5 through d.i.v. 9, that encompasses the transition of GABAergic cells from excitatory to inhibitory (*11, 40, 41*). In WT GAD65(−) glutamatergic neurons, the integrated area (**Figure 3A-B**) and peak amplitude (**Figure 3A** and **3C**) of the evoked α7 nAChR Ca^2+^ transients sharply increased from d.i.v. 5 to d.i.v. 7 and remained elevated at d.i.v. 9. In contrast, *Fmr1*^KO^ GAD65(−) neurons did not exhibit a significant increase in the integrated area and peak amplitude of the response at d.i.v. 7 compared to d.i.v. 5 (**Figure 3A-C**), suggesting a deviation in the typical progression of the α7 nAChR response in *Fmr1*^KO^ neurons. Notably, the α7 nAChR responses in *Fmr1*^KO^ glutamatergic neurons were significantly weaker compared to WT, as shown by reduced mean integrated area (**Figure 3D**) and peak amplitude (**Figure 3E**) at d.i.v. 7 and d.i.v. 9.

**Figure 3.**
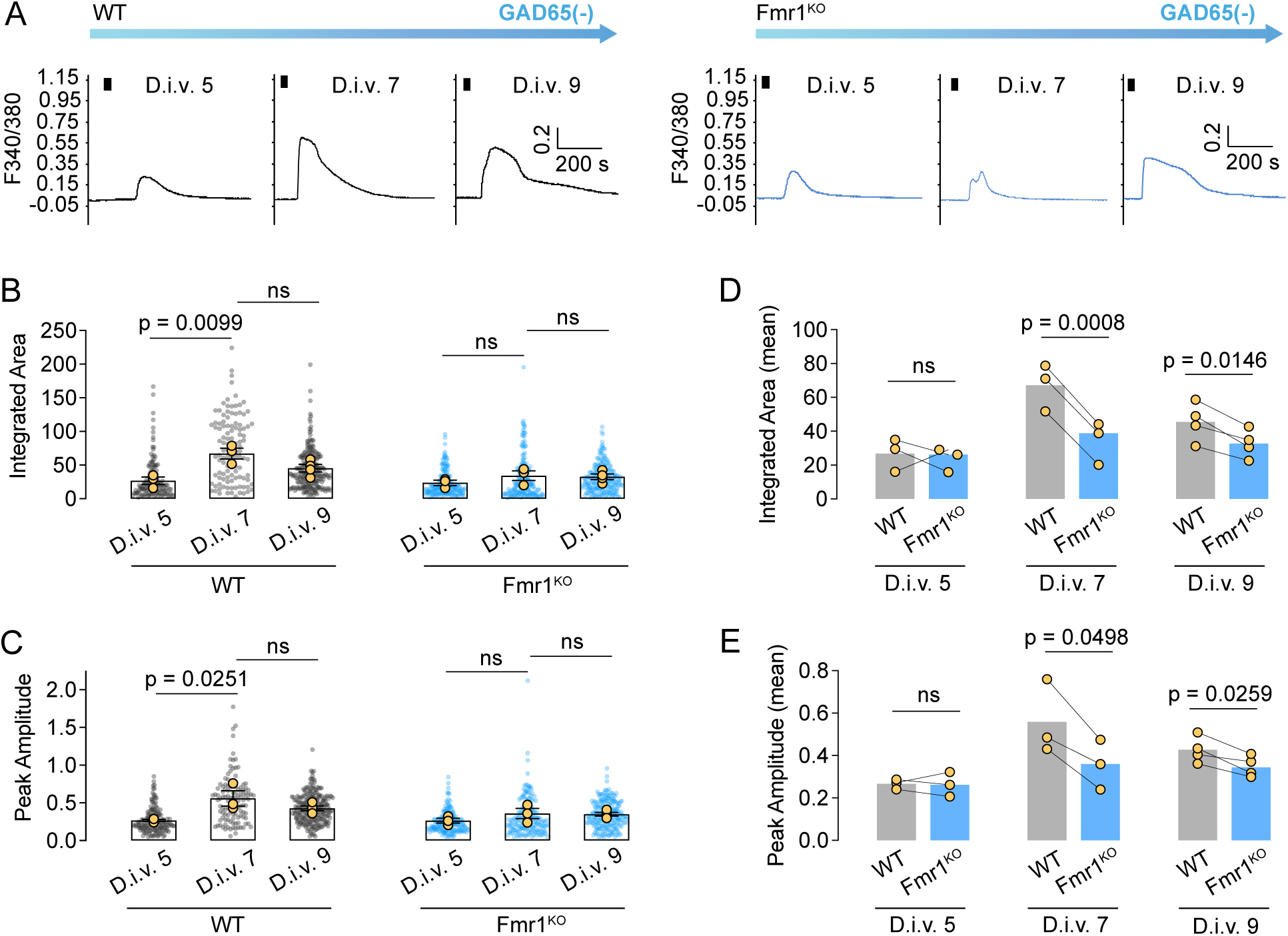
Agonist-evoked α7 nAChR Ca^2+^ responses are dampened in *Fmr1*^KO^ glutamatergic neurons during early neuronal maturation. **(A)** Representative traces of Ca^2+^ responses evoked by 100 nM PNU-282987 with 3 µM PNU-120596 (black bars) in WT (left) and *Fmr1*^KO^ (right) GAD65(−) neurons during maturation *in vitro* at d.i.v. 5, 7, 9. (**B)** Quantification of the integrated area of the responses. Yellow symbols are the means of individual experiments (d.i.v. 5-7 N = 3, d.i.v. 9 N = 4), filled symbols are all measured responses (WT d.i.v. 5 n = 139, d.i.v. 7 n = 112, d.i.v. 9 n = 231; *Fmr1*^KO^ d.i.v. 5 n = 172, d.i.v. 7 n = 170, d.i.v. 9 n = 216), one way ANOVA with Tukey’s post-test per group, WT *p* = 0.0123, *Fmr1*^KO^ *p* = 0.3743. (**C**) Quantification of peak amplitude over time in GAD65(−) neurons. Yellow symbols are means (d.i.v. 5-7 N = 3, d.i.v. 9 N = 4), filled symbols are all measured responses (WT d.i.v. 5 n = 148, d.i.v. 7 n = 115, d.i.v. 9 n = 232; *Fmr1*^KO^ d.i.v. 5 n = 176, d.i.v. 7 n = 180, d.i.v. 9 n = 219); one way ANOVA with Tukey’s post-test per group, WT *p* = 0.0302, *Fmr1*^KO^ *p* = 0.3018. (**D**) Paired comparison of WT and *Fmr1*^KO^ mean integrated area of Ca^2+^ responses in GAD65(−) neurons in matched cultures at different ages *in vitro*. Mean ± SEM, d.i.v. 5 WT 26.8 ± 5.63, KO 23.7 ± 4.0 N = 3; d.i.v. 7 WT 67.2 ± 8.04, KO 34.4 ± 7.26 N = 3; d.i.v. 9 WT 45.5 ± 5.73, KO 32.7 ± 4.21 N = 4, paired t-test. (**E**) Paired comparison of WT and *Fmr1*^KO^ mean peak amplitude of Ca^2+^ responses in GAD65(−) neurons in matched cultures at different ages *in vitro*. Mean ± SEM, d.i.v. 5 WT 0.27 ± 0.014, KO 0.26 ± 0.033 N = 3; d.i.v. 7 WT 0.56 ± 0.1, KO 0.36 ± 0.068 N = 3; d.i.v. 9 WT 0.43 ± 0.031, KO 0.35 ± 0.025 N = 4; paired t-test.

Similarly to glutamatergic neurons, in WT GAD65(+) GABAergic neurons the evoked α7 nAChR Ca^2+^ transients were potentiated at d.i.v. 7 relative to d.i.v. 5 but declined by d.i.v. 9, as shown by the integrated area of the response (**Figure 4A-B**). *Fmr1*^KO^ GAD65(+) neurons did not exhibit a transient potentiation of α7 nAChR responses at d.i.v. 7 (**Figure 4A-C**) but instead showed a marked reduction in the integrated area compared to WT at d.i.v. 7 and d.i.v. 9 (**Figure 4E**). Although the peak amplitude of the evoked responses did not differ significantly between genotypes (**Figure 4A-C**) we noted a significant correlation between the integrated area and the extended duration of the response measured by a Response Decline Index (**Figure 4D**) (Spearman r –0.4709, n = 54 cells, *p* = 0.0003) calculated as the ratio of the integrated area for the first 100 s (typically encompassing the rise to peak) over the entire duration of the response back to baseline. The reduction in integrated area in the *Fmr1*^KO^ was not driven by a difference in peak amplitude (**Figure 4F**) but rather by a reduced duration of the response (**Figure 4G**). Altogether, these data point to a stalled development of α7 nAChR Ca^2+^ responses in *Fmr1*^KO^ glutamatergic and GABAergic neurons and an overall dampening of α7 nAChR activity compared to WT during their maturation.

**Figure 4.**
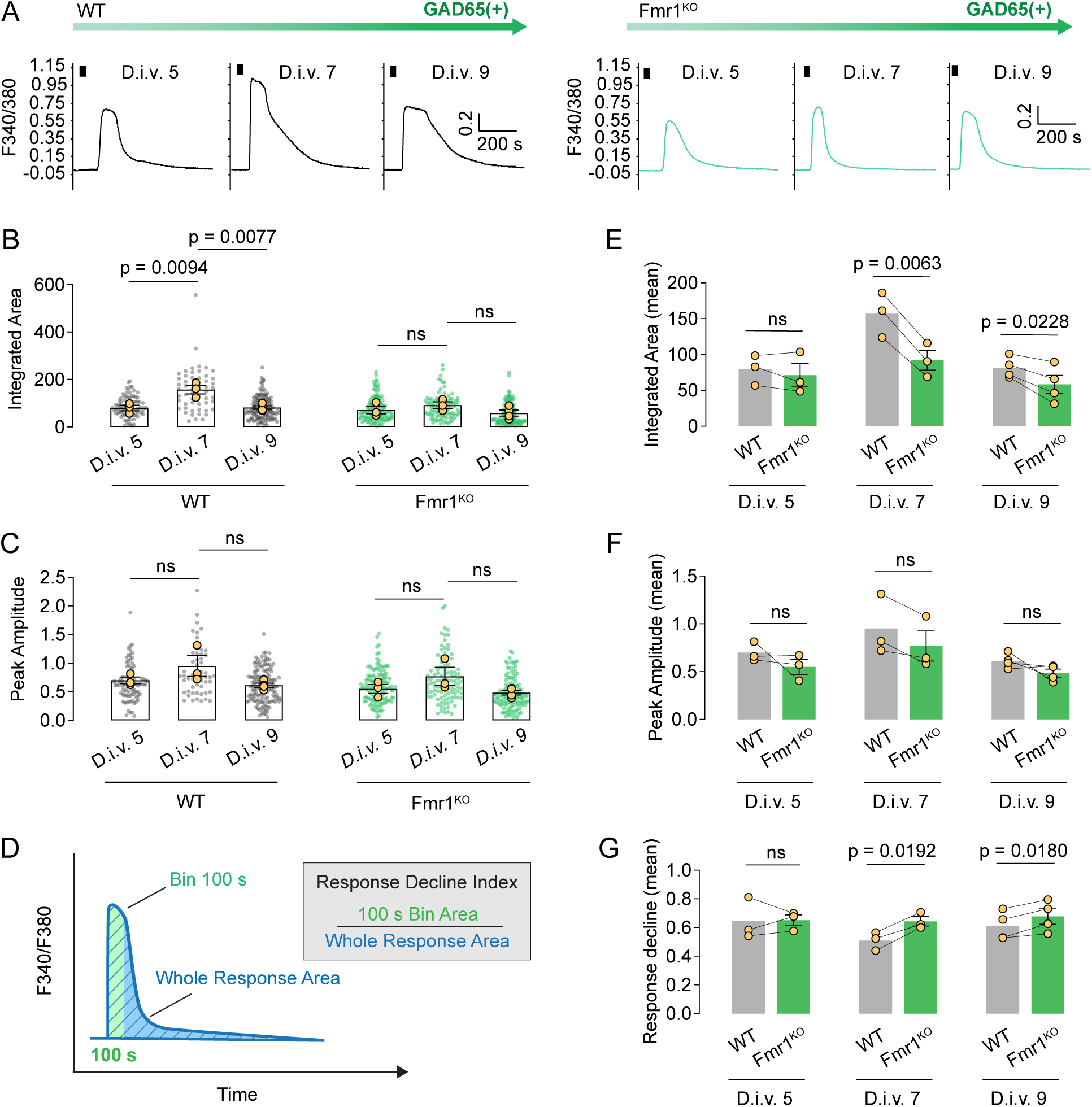
Agonist-evoked α7 nAChR Ca^2+^ responses are dampened in *Fmr1*^KO^ GABAergic cells during early neuronal maturation. **(A)** Representative traces of Ca^2+^ responses evoked by 100 nM PNU-282987 with 3 µM PNU-120596 (black bars) in WT (left) and *Fmr1*^KO^ (right) GAD65(+) cells during maturation *in vitro* at d.iv. 5, 7, 9. (**B)** Quantification of the integrated area in WT and *Fmr1*^KO^ GAD65(+) neurons. Yellow symbols are means of individual experiments (d.i.v. 5-7 N = 3, d.i.v. 9 N = 4), filled symbols are all measured responses (WT, d.i.v. 5 n = 85, d.i.v. 7 n = 54, d.i.v. 9 n = 135; *Fmr1*^KO^ d.i.v. 5 n = 126, d.i.v. 7 n = 99, d.i.v. 9 n = 116); one way ANOVA with Tukey’s post-test per group, WT *p* = 0.0053, KO *p* = 0.2982. (**C**) Quantification of peak amplitude in WT and *Fmr1*^KO^ GAD65(+) neurons. Yellow symbols are means (d.i.v. 5-7 N = 3, d.i.v. 9 N = 4); filled symbols are all measured responses (WT, d.i.v. 5 n = 87, d.i.v. 7 n = 54, d.i.v. 9 n = 136; *Fmr1*^KO^ d.i.v. 5 n = 128, d.i.v. 7 n = 101, d.i.v. 9 n = 117); one way ANOVA with Tukey’s post-test per group, WT *p* = 0.1168, KO *p* = 0.1563. (**D**) Schematic of Response Decline Index to measure response duration. (**E**) Paired comparison of WT and *Fmr1*^KO^ mean integrated area in GAD65(+) neurons in matched cultures at different ages *in vitro*. Mean ± SEM, d.i.v. 5 WT 79.5 ± 12.1, KO 71.3 ± 16.6 N = 3; d.i.v. 7 WT 157 ± 18.1, KO 91.9 ± 13.6 N = 3; d.i.v. 9 WT 81.7 ± 7.42, KO 58.4 ± 12.6 N = 4, paired t-test. (**F**) Paired comparison of WT and *Fmr1*^KO^ mean peak amplitude in GAD65(+) neurons in matched cultures at different ages *in vitro*. Mean ± SEM, d.i.v. 5 WT 0.7 ± 0.059 KO 0.55 ± 0.078 N = 3, d.i.v. 7 WT 0.95 ± 0.18 KO 0.77 ± 0.16 N = 3, d.i.v. 9 WT 0.61 ± 0.038 KO 0.49 ± 0.042 N = 4; paired t-test. (**G**) Paired comparison of Response Decline Index in matched cultures. Mean ± SEM, d.i.v. 5 WT 0.65 ± 0.084, KO 0.65 ± 0.038 N = 3, d.i.v. 7 WT 0.51 ± 0.037, KO 0.64 ± 0.032 N = 3, d.i.v. 9 WT 0.61 ± 0.05, KO 0.68 ± 0.053 N = 4; paired t-test.

### Reducing Ly6H expression enhances α7 nAChR Ca^2+^ responses in *Fmr1*^KO^ neurons

Having found that the evoked α7 nAChR-dependent Ca^2+^ response is dampened in *Fmr1*^KO^ neurons, we considered the possibility that expression of the α7 subunit might be altered in the *Fmr1*^KO^ given that FMRP can regulate mRNA stability and translation (*5*). As determined by Western blot, the total abundance of α7 in hippocampal neurons at d.i.v. 5, 7 and 9 did not significantly differ between WT and *Fmr1*^KO^ (**Figure S4A-B**). In addition, labeling intact hippocampal neurons with fluorescent α-Bungarotoxin (αBGTX), which binds selectively and with high affinity to the mature α7 nAChR homopentamer, did not reveal significant differences in the density of surface bound αBGTX in WT and *Fmr1*^KO^ (**Figure S4C-D**). Congruent with its expression profile in primary hippocampal neurons, α7 expression in early postnatal hippocampus (P8) was comparable between genotypes (**Figure S4E-F**), altogether suggesting that dampened α7 nAChR responses in the *Fmr1*^KO^ are unlikely to arise from deficits in α7 expression.

As previously reported, Ly6H can restrain α7 nAChR activity by directly binding to α7 nAChRs inserted in the plasma membrane (*29*). The reported inhibition of α7 nAChR function by Ly6H is analogous to the action of Lynx1, the first member of the Ly6/uPAR family shown to bind to and inhibit nAChRs, operating as an endogenous “brake” for nAChR signaling (*25*). Because in *Fmr1*^KO^ neurons the decrease in evoked α7 nAChR responses is concomitant with increased surface expression of Ly6H, we considered the possibility that reducing Ly6H expression may potentiate α7 nAChR activity in isolated neurons. To test this possibility, we infected *Fmr1*^KO^ hippocampal neurons at d.i.v. 2 with viruses encoding eGFP together with a previously tested Ly6H-specific shRNA or with scrambled control shRNA (*30*) and examined them at d.i.v. 9. Western blot analysis of neurons treated with Ly6H shRNA confirmed efficient downregulation of the endogenous protein compared to control shRNA-treated cells (**Figure S5A-B**). Evoked α7 nAChR Ca^2+^ responses were induced by bath-application of agonist PNU-282987 together with the PAM PNU-120596, as described above, in eGFP-positive neurons that were immunolabeled post-hoc for Ly6H and GABA expression (**Figure 5A**) to confirm downregulation and distinguish GABAergic (GABA+) *versus* glutamatergic neurons (GABA-), respectively. In both GABA(−) (**Figure 5A-C**) and GABA(+) neurons (**Figure 5A** and **5D-E**) Ly6H knockdown resulted in α7 nAChR Ca^2+^ responses with larger integrated areas compared to neurons that received scrambled shRNA. Thus, lowering the expression of Ly6H can potentiate the dampened α7 nAChR Ca^2+^ responses in *Fmr1*^KO^ neurons.

**Figure 5.**
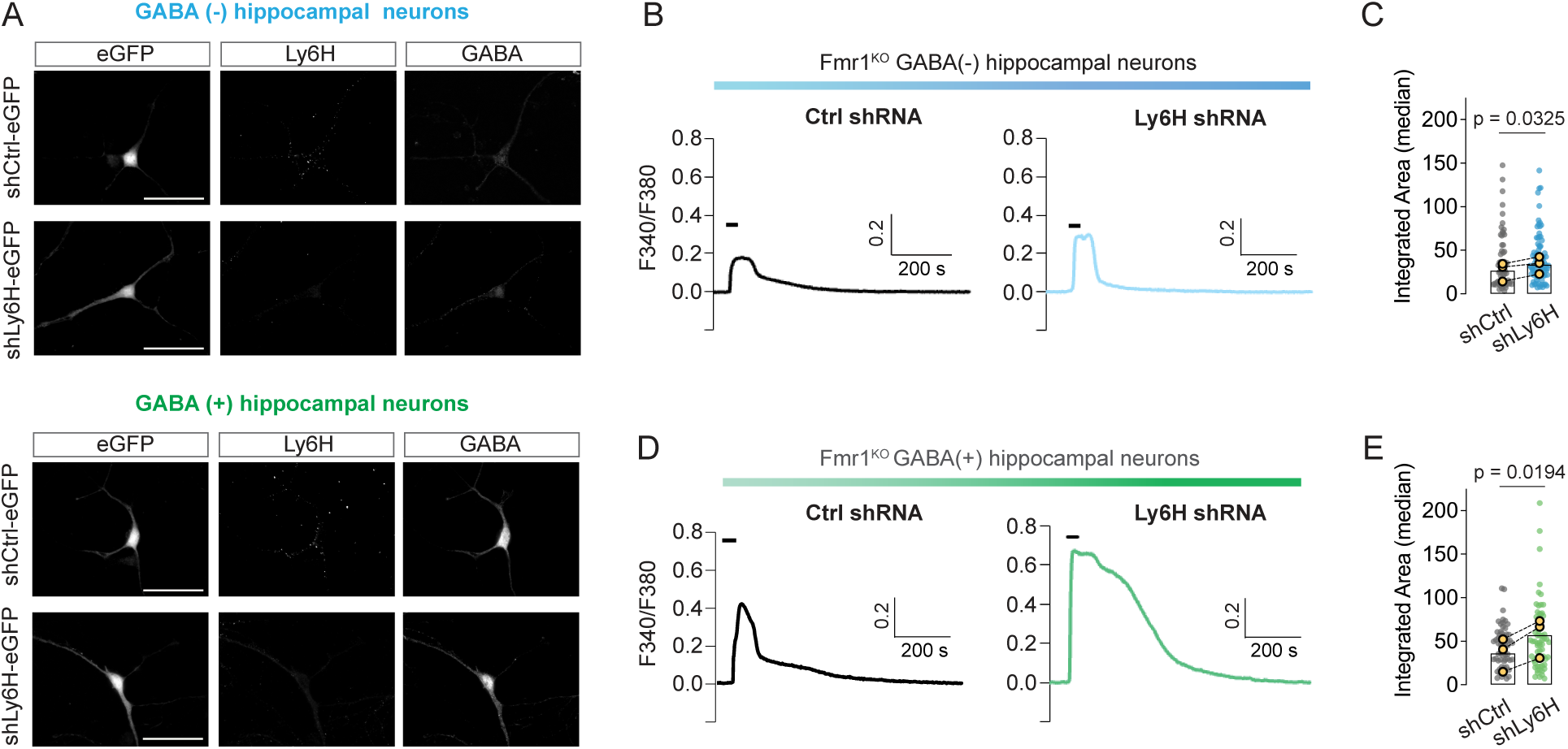
Ly6H knockdown in *Fmr1*^KO^ hippocampal neurons potentiates evoked α7 nAChR Ca^2+^ responses. **(A)** Representative images of d.i.v. 9 *Fmr1*^KO^ hippocampal neurons transduced with Ly6H shRNA (shLy6H) or control scrambled shRNA (shCtrl) co-expressing eGFP. Cells were immunolabeled for Ly6H and GABA after Ca^2+^ imaging; scale bars, 50 µm. (**B**) Representative Ca^2+^ responses induced by 100 nM PNU-282987 with 3 µM PNU-120596 (black bars) in d.i.v. 9 GABA(−) neurons transduced with shLy6H or shCtrl. **(C)** Quantification of the integrated area of the response in GABA(−) neurons; filled symbols are all measured responses (shLy6H n = 86), shCtrl n = 60), yellow symbols are the medians of N = 3 experiments; paired t-test. (**D**) Representative Ca^2+^ responses evoked as in (B) in d.i.v. 9 GABA(+) *Fmr1*^KO^ neurons transduced with shLy6H or shCtrl. (**E**) Quantification of integrated area in GABA(+) neurons; filled symbols are all measured responses (shLy6H n = 64, shCtrl n = 60), yellow symbols are the medians of N = 3 experiments; paired t-test.

### Boosting α7-nAChR activity by systemic administration of the PAM PNU-120596 improves behavioral deficits in *Fmr1*^KO^ mice

*Fmr1*^KO^ mice exhibit abnormal hippocampal synaptic plasticity, including long-term potentiation and long-term depression, supporting the notion that synaptic disturbances underlie learning and memory deficits associated with FXS (*42*). Accordingly, *Fmr1*^KO^ mice exhibit deficits in hippocampal-based memory tasks including object location memory (OLM) (*43–45*). OLM is a form of spontaneous recognition memory that emerges early in mouse development (*46*), is typically sensitive to cholinergic activity (*43*), and is lost after administration of methyllycaconitine or deletion of *Chrna7* in interneurons (*47*). Because of the intrinsic α7-nAChR hypofunction observed in *Fmr1*^KO^ hippocampal neurons, we tested whether pharmacological enhancement of α7-nAChR activity in *Fmr1*^KO^ mice could ameliorate deficits in hippocampal-based memory. Pilot studies with naïve WT and *Fmr1*^KO^ P22 mice tested for their ability to distinguish the novel location of an object 24 h after a training phase (24 h retention interval) (**Figure S6A**) confirmed significant deficits in exploratory preference for a novel location in early adolescent *Fmr1*^KO^ mice compared to WT (**Figure S6B-D**). To enhance α7 nAChR activity, *Fmr1*^KO^ mice received daily intraperitoneal injections (i.p.) of the α7 nAChR-selective PAM PNU-120596 (3 mg/kg) or vehicle from P18 to P22 (test day), whereas WT mice received vehicle only (**Figure 6A-B**). Although the total time spent with objects did not differ across genotype and treatment conditions (**Figure 6C**), vehicle-treated *Fmr1*^KO^ mice exhibited reduced exploratory preference for objects in the novel location compared to both WT mice treated with vehicle and *Fmr1*^KO^ mice treated with PNU-120596 (**Figure 6B** and **6D**), indicating that α7 nAChR enhancement improves spatial memory in mutant mice.

**Figure 6.**
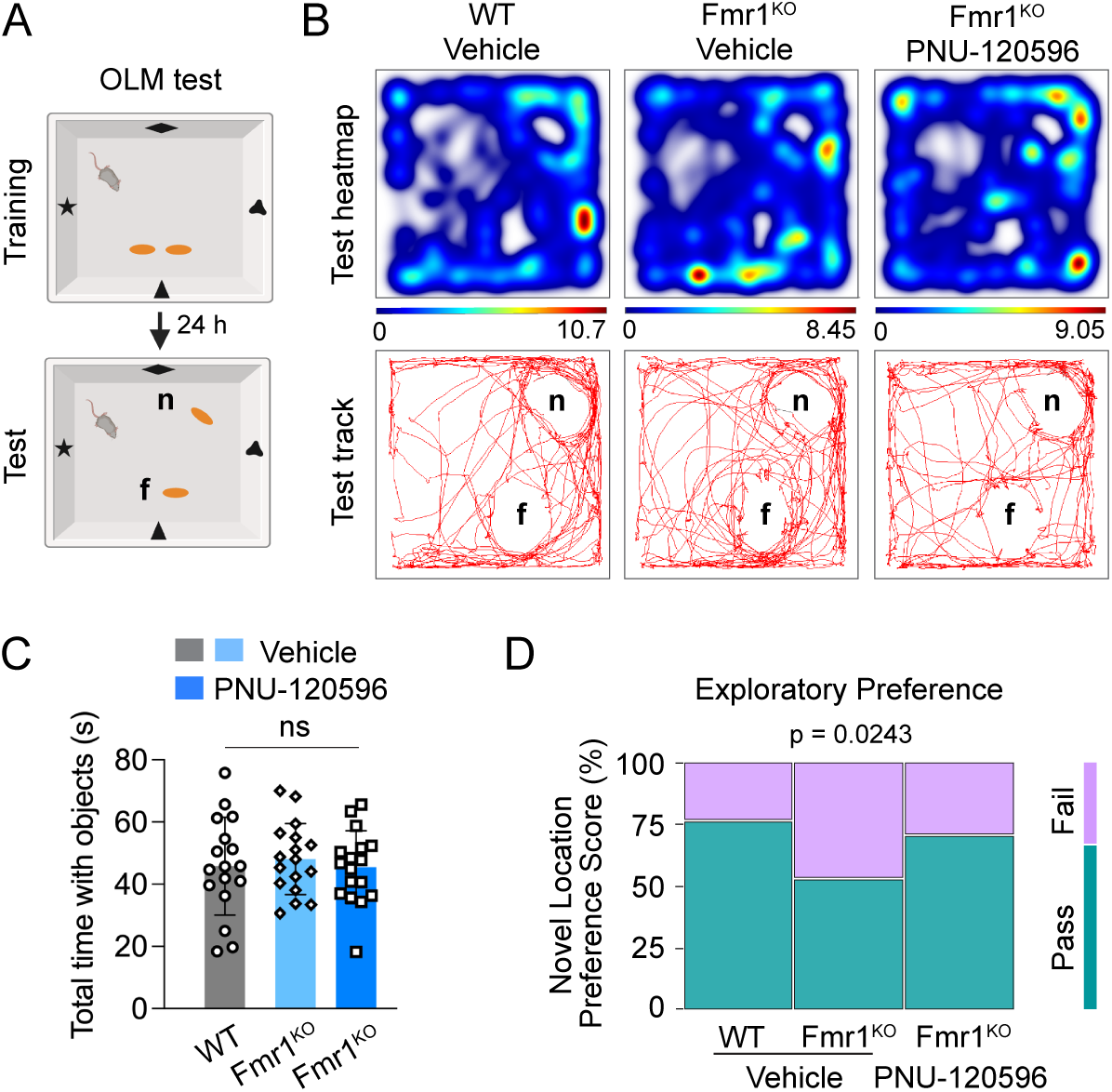
The α7 nAChR positive allosteric modulator PNU-120596 improves spatial memory in *Fmr1*^KO^ mice. **(A)** Schematic of OLM test where *n* indicates the novel location and *f* the familiar object location (created with BioRender). (**B**) Representative activity heatmaps and tracks of mice during the test phase (5 min) of the OLM test. (**C**) Quantification of total time spent with objects by WT mice treated with vehicle and *Fmr1*^KO^ mice treated with vehicle or PNU-120596 (n = 17 per group); one way ANOVA, *p* = 0.8191. (**D**) Mosaic plot of Exploratory Preference Score expressed as Z-score stratified by sex, where Pass (preference for novel location) corresponds to Z ≥ –1 (n = 17 mice per group); Fisher’s Exact Test (3×2).

In addition to cognitive deficits, individuals with FXS frequently manifest hyperactivity, are commonly affected by disabling auditory hypersensitivity, and experience seizures. Both hyperactivity and auditory hypersensitivity in the form of audiogenic seizures (AGS) have been reproducibly reported to occur in *Fmr1*^KO^ mice, with susceptibility to AGS frequently used in preclinical drug studies of FXS (*48, 49*). Notably, compromised α7 nAChR signaling has been typically implicated in the development of hyperactivity and seizures in animal models (*50–53*) and human subjects (*18, 54*). Given the cell intrinsic nature of α7 nAChR hypofunction in the *Fmr1*^KO^, we hypothesized that enhancing α7 nAChR activity could have broad impact on circuit excitability and improve the hyperlocomotion exhibited by early adolescent *Fmr1*^KO^ mice (**Figure S7**) and AGS outcomes. To test these possibilities, locomotor activity and AGS were sequentially assessed (in that order) in P22 *Fmr1*^KO^ mice treated for five days with PNU-120596 or vehicle (**Figure 7A**). Compared to the vehicle-treated, *Fmr1*^KO^ mice treated with PNU-120596 exhibited a decrease in the total distance traveled (**Figure 7B** and **7C**), duration of movement (**Figure 7D**) and frequency of movement initiation (**Figure 7E**). Next, mice were exposed to a loud alarm (∼120 dB) for one minute to trigger AGS initiation. AGS are generalized convulsive motor seizures triggered by acoustic stimulation that progress in several phases, starting from an initial period of wild running which can be followed by a tonic-clonic seizure and respiratory failure/cardiac arrest depending on seizure severity (**Figure S8A**). In *Fmr1*^KO^ mice, AGS are reproducibly observed across ages and sexes with P22 being the most susceptible age (*48*). The progression and severity of AGS were scored according to an established metric, the Audiogenic Response Score (ARS) (**Figure S8B**), that categorizes seizure severity from 1 to 9, with ARS 1 representing the least severe (wild running only) (*55*). Administration of PNU-120596 reduced the severity of AGS compared to vehicle as indicated by less extreme ARS scores. Interestingly, a larger proportion of mice treated with PNU-120596 experienced two successive bouts of wild running (WR2) (**Figure 7F**). The appearance of WR2 before presentation of a tonic-clonic phase indicates less severe seizure compared to one bout followed by a seizure, because longer stimulation is required to set off seizure progression (*56, 57*). In aggregate, PNU-120596 administration resulted in less severe ARS scores, with a significant decrease in the percentage of mice that received an ARS score of 9 (**Figure 7G**) and a higher percentage of mice with an ARS score of 4 and 8. Importantly, PNU-120596 reduced mortality following seizures (**Figure 7H**). Altogether, these results indicate that treatment with an α7 nAChR-selective PAM to counter α7 nAChR hypofunction in *Fmr1*^KO^ mice, ameliorates memory impairment and reduces hyperactivity and seizure severity.

**Figure 7.**
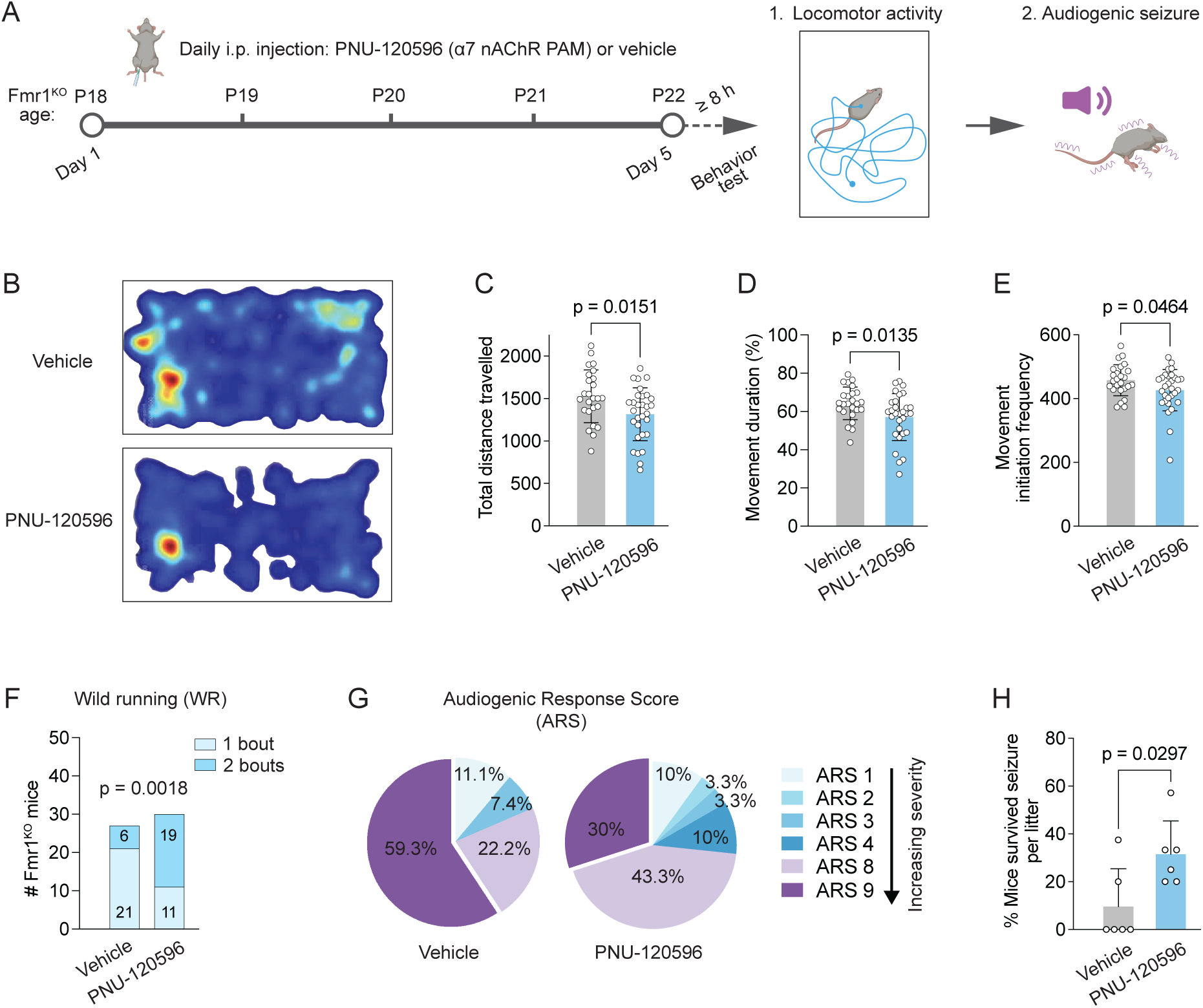
The α7 nAChR positive allosteric modulator PNU-120596 decreases hyperactivity and seizure severity in *Fmr1*^KO^ mice. **(A)** Schematic of experimental strategy and timeline (created with BioRender). **(B)** Representative heatmaps of activity in open field of *Fmr1*^KO^ mice treated with PNU-120596 or vehicle. **(C)** Total distance traveled; mean ± SEM, vehicle 1527 ± 62 cm n = 25, PNU-120596 1315 ± 57 n = 30 mice. (**D**) Percentage of time spent in movement; mean ± SEM, vehicle 64 ± 1.7 n = 25, PNU-120596 57 ± 2.2 n = 30. (**E**) Frequency of movement initiation; mean ± SEM, vehicle 458 ± 9.7 n = 25, PNU-120596 426 ± 12 n = 30, unpaired t-test with Welch correction. (**F**) Analysis of audiogenic seizure severity. Number of *Fmr1*^KO^ mice exhibiting one bout of wild running (WR1) *versus* two (WR2); vehicle n = 27 mice tested, PNU-120596 n = 30; two-sided Chi-square. (**G**) Percentage of *Fmr1*^KO^ mice treated with vehicle (n = 27) or PNU-120596 (n = 30) scoring in respective Audiogenic Response Score (ARS) category; only observed ARS categories are displayed. A greater number of vehicle-treated mice experienced seizures with ARS9 *vs*. 8, compared to mice treated with PNU-120596: *p* = 0.0331, two-sided Chi-square. (**H**) Percentage per litter of vehicle and PNU-120596 treated *Fmr1*^KO^ mice that survived after seizure; mean ± SEM, vehicle 9.6 ± 6.5, PNU-120596 31 ± 5.7 n = 6 litters, unpaired t-test with Welch correction.

## DISCUSSION

Here, we report the identification of cell-intrinsic abnormalities in the α7 nAChR signaling system in a mouse model of FXS, and present evidence that their rescue ameliorates cognitive deficits and behavioral pathologies linked to hyperexcitability. Our results show that the trafficking of Ly6H, an α7 nAChR auxiliary protein, is altered in hippocampal *Fmr1*^KO^ neurons, resulting in its increased surface expression. Concomitantly, evoked α7 nAChR Ca^2+^ responses are dampened in *Fmr1*^KO^ neurons of glutamatergic and GABAergic lineage in absence of detectable differences in α7 expression. α7 nAChRs hypofunction in *Fmr1*^KO^ neurons is relieved by reducing Ly6H abundance, pointing to cell-intrinsic deficits in α7 nAChR responses hinging on mis-expression of Ly6H. Congruent with deficiency in α7 nAChR activity, *in vivo* administration of the bioactive positive allosteric modulator PNU-120596 – which increases evoked α7 nAChR-mediated Ca^2+^ influx – ameliorates spatial memory deficits, hyperlocomotion and seizure severity in *Fmr1*^KO^ mice.

It is firmly established that both the expression of functional nAChRs and the strength and duration of their responses are the product of multiple highly regulated cellular processes, encompassing controlled biogenesis (*e.g*. folding, assembly) and export, and on-site modification of receptor properties. These processes are coordinated through cell-specific engagement of selected chaperones and auxiliary proteins and, therefore, dependent on their respective temporal and spatial profiles of expression. The identification of these critical components of the “nAChR system” has provided key insights towards a more complete understanding of nAChR properties in pathophysiological conditions (*22*). Our results highlight Ly6H as one such critical component that could contribute to deficits in α7 nAChR function in *Fmr1*^KO^ neurons. Mechanistically, the observed alterations in Ly6H surface expression and localization may arise from aberrant processing of its GPI anchor and or abnormalities in Golgi-plasma membrane transport. Indeed, several enzymes involved in the processing of GPI-anchored proteins are encoded by FMRP-client mRNAs (*58, 59*). In heterologous expression systems, Ly6H overexpression reduces α7 nAChR export to the surface and agonist binding, overall resulting in reduced evoked responses (*26*). Conversely, downregulation of endogenous Ly6H was shown to boost α7 nAChR activity in rat hippocampal neurons (*26, 27*). Importantly, Ly6H can curb α7 nAChR responses by directly binding to the receptor itself when expressed in the plasma membrane, an effect abrogated by cleavage of the Ly6H GPI anchor or by mutations in the 3-finger domain required for nAChR binding and structurally homologous to that of αBGTX (*29*). CNS-expressed α7 nAChRs desensitize very rapidly and undergo prolonged residual inhibition (*33*). Several Ly6/uPAR family proteins have been shown to regulate nAChR desensitization either by accelerating (*e.g*. Lynx1, Lynx2) (*25, 60*) or delaying it (Ly6g6e) (*27*) resulting in net dampening or potentiation of the response, respectively. Endogenous Ly6H may reduce α7 nAChR activity in a membrane-delimited manner by promoting desensitization like Lynx1/2 proteins, so that abnormal crowding of Ly6H in somatodendritic regions of *Fmr1*^KO^ neurons could lead to locally depressed α7 nAChR responses. Our experimental results provide some support for this possibility since we did not detect reductions in α7 nAChR expression in *Fmr1*^KO^ neurons that could account for the dampened evoked responses. Crucially, downregulation of Ly6H expression was effective in boosting α7 nAChR activity.

Our results show a robust, but transient upregulation of α7 nAChR evoked activity during the early maturation of WT hippocampal neurons *in vitro,* prior to the formation of excitatory synaptic contacts and the full maturation of interneurons. Notably, the transient increase in α7 nAChR activity – occurring at a time of heightened Ly6H mis-expression in mutant neurons – was compromised in the *Fmr1*^KO^. Since the expression level of α7 subunits remained stable over time in both genotypes, these results suggest that the observed differences in the strength of α7 nAChR responses may depend on the regulated expression of α7 nAChR-associated proteins. Multiple members of the Ly6/uPAR family are expressed in hippocampal neurons and their expression is developmentally regulated (*24, 34, 61*). Thus, potential competition between Ly6/uPAR proteins with opposing function (*e.g.* increasing *vs.* decreasing nAChR desensitization) may contribute to shaping the developmental progression of α7 nAChR responses. Aberrant expression of Ly6H mRNA was observed in FXS patient-derived iPSCs differentiated into dorsal forebrain neurons of glutamatergic lineage (*62*). Notably, Ly6H was found to be one of the most upregulated mRNAs in human FXS glutamatergic lineage neurons at late stages of *in vitro* differentiation, whereas it was downregulated at early stages. Interestingly, an independent study in FXS iPSC-derived immature neurons reported downregulation of Lynx1 (*63*). Thus, dysfunctions in nAChR auxiliary proteins that impact the maturation of functional nAChR responses are present in *Fmr1*^KO^ mice and FXS patients alike.

In the hippocampus, α7-nAChR Ca^2+^ signaling contributes to the maturation of excitatory and inhibitory neurons (*11, 13, 64*) and synapses (*10, 12*), as well as circuit formation and function during early postnatal development (*7, 11, 65*). The significantly more robust α7 nAChR responses that we experimentally observed in immature GABAergic cells suggest that α7 nAChRs could play a larger role in shaping inhibition, congruent with evidence of inhibitory deficits and interneuron dysfunction in FXS (*66*). Deficits in α7 nAChR signaling in *Fmr1*^KO^ neurons could be particularly impactful during critical periods of development when synaptic connectivity and neuronal networks are being formed and stabilized. Spatial memory, exemplified by the ability to recognize the novel location of an object, was shown to emerge in mice around P21 reflecting the ontogeny of hippocampal function development (*67, 68*) and is dependent on α7 nAChR activity (*47*). Our results show that the capacity for long-term (24 h) retention of spatial information in *Fmr1*^KO^ mice is improved by early and sustained potentiation of α7 nAChRs, supporting its pro-cognitive impact in the mutant during a critical developmental period.

nAChR auxiliary proteins such as Ly6H, Lynx1/2 and Lypd6 can associate with other nAChRs including α4β2 nAChR, which is highly expressed in cortical regions. Thus, in addition to α7, other nAChRs may exhibit intrinsically altered activity in *Fmr1*^KO^ mice in a regional and cell-type specific manner. Our results indicate that aberrant trafficking of Ly6H and intrinsic deficits in α7 nAChR responses occur early during hippocampal development (P7) and neuronal maturation, respectively, ages that are relevant to FXS clinical manifestations (*69*). Although most studies in *Fmr1*^KO^ mice have used juvenile or adult animals, recent evidence supports the notion that some cellular and synaptic pathologies in mutant mice may be compensatory and arise to mitigate primary deficits that originated during development (*70*). Related to this, enhanced cholinergic tone was reported in adult *Fmr1*^KO^ mice and linked to impaired attentional behavior (*71*). Thus, it will be important to define the temporal dynamics of expression of auxiliary proteins and functional maturation of nAChRs in FXS, that might differ across brain regions and cell-types.

Although our cellular studies focused on hippocampal neurons, both α7 and Ly6H are widely expressed in the CNS suggesting that intrinsic dysfunctions in α7 nAChR activity might be present in different brain regions. α7 nAChRs participate in many of the processes that go awry in individuals with FXS including attention, hyperactivity, and seizures. About ∼80% of FXS patients are diagnosed with attention-deficit hyperactivity disorder and, likewise, *Fmr1*^KO^ mice exhibit locomotor hyperactivity linked to circuit hyperexcitability. Several studies have reported early network hyperexcitability in *Fmr1*^KO^ mice that may underlie seizure susceptibility (*72*). *Fmr1*^KO^ mice are highly susceptible to AGS – considered an expression of sensory hypersensitivity – that manifest around P17 and peak at approximately P22, persisting into adulthood (*48*). We find that administration of PNU-120596 ameliorates hyperlocomotion in *Fmr1*^KO^ mice, in agreement with previous work demonstrating that deficits in α7 nAChR signaling are causally related to the manifestation of hyperactivity. Moreover, our results indicate that timed and sustained delivery of PNU-120596 is effective in reducing AGS severity at P22 when their manifestation is most severe. PNU-120596 may delay seizure progression or change seizure threshold by promoting the brain’s natural dampening mechanisms.

Alterations in muscarinic ACh receptor signaling and expression were also reported in FXS. Interestingly, despite upregulated expression of M1 and M4 receptors in *Fmr1*^KO^ mice, pharmacological enhancement of M4 activity with a PAM was found to be beneficial for amelioration of audiogenic seizures, suggesting rescue of latent hypofunction (*15*). Our results, together with the above-mentioned findings in *Fmr1*^KO^ mice and patients’ derived cells, converge to suggest a broad cholinergic deficit in FXS. Deficits in cholinergic signaling have been reported in other neurodevelopmental disorders, such as Rett syndrome, both in patient-derived neurons and a mouse model of the condition (*50, 73*). Moreover, cholinergic enhancement was shown to improve cognitive flexibility and social interaction in mouse models of autism and, congruent with preclinical studies, small open-label trials identified beneficial effects of targeting nAChRs in autism patients (*74*). We propose that our preclinical findings support the validity of pursuing α7-based intervention in FXS. A primary need for this approach is the availability of safe and effective drugs. PNU-120596 is a type II PAM that inhibits α7 nAChR desensitization and enhances the potency of nicotinic agonists without activation of the receptors when administered alone. It is specific for α7 nAChRs, with no detectable effect on α4β2, α3β4 and α9α10 nAChRs, and has been shown to be bioactive and effective in many studies in rodents in which it is well tolerated. However, safety concerns because of potential excessive increase in intracellular Ca^2+^ have prevented clinical testing. The type I PAM AVL-3288 could be considered an alternative since it has been successfully evaluated for safety and neurocognitive effects in healthy human subjects (*75*). Type I PAMs enhance α7 nAChR cholinergic activation but preserve its rapid desensitization, alleviating concerns of excessive Ca^2+^ influx. To move forward, preclinical studies could first establish dose and effectiveness of AVL-3288 in ameliorating hyperexcitability-related pathologies in *Fmr1*^KO^ mice and define the timing of interventions. A critical point would be to determine whether administration during a narrow temporal window early in development (*e.g.* P4-P8) could be sufficient for correction of deficits during adolescence or whether chronic or repeated treatment are instead needed. Nevertheless, given the lifelong persistence of FXS associated pathologies that are linked to α7 nAChR hypofunction, interventions with α7 PAMs could also be beneficial in adult subjects.

## MATERIALS AND METHODS

### Animals

WT and *Fmr1*^KO^ (FVB.129P2-*Pde6b*^+^ strain) mice were obtained from The Jackson Laboratories (Bar Harbor, ME) and bred in-house. Mice were fed ad libitum and housed with a 12 h light/dark cycle. All animal procedures were carried out according to protocols approved by the Albert Einstein College of Medicine, in accordance with the Guide for the care and use of laboratory animals by the United States PHS.

### Genotyping

Genomic DNA was prepared by tissue digestion in a buffer of 50 mM NaCl, 10 mM Tris-HCl pH.9, 0.4% NP-40, 0.4% Tween-20 and Proteinase K (New England Biolabs) for 1 h at 55°C followed by incubation at 95°C for 12 min and centrifugation at 5000 rpm for 12 min. Oligonucleotides used were: GTGGTTAGCTAAAGTGAGGATGAT (forward; oligo.1) and GTGGGCTCTATGGCTTCTGAGG (reverse; oligo.2) for amplification of the *Fmr1*^KO^ allele and oligo.1 together with CAGGTTTGTTGGGATTAACAGATC (reverse; oligo.3) for the WT allele, respectively. Annealing temperature was 56°C for WT and 60°C for *Fmr1*^KO^ amplification, with extension temperature 68°C for WT and 72°C for *Fmr1*^KO^, cycled 35 times.

### Primary hippocampal cultures

Primary cultures were prepared from P0/P1 pups generated by crossing *Fmr1* heterozygous females with WT males. The brains were stored in the dark at 4°C in Hibernate A medium (Gibco) until genotype determination. Hippocampi were microdissected in Ca^2+/^Mg^2+^-free Hanks Balanced Saline Solution (HBSS), finely chopped, and incubated for 30 min at 37°C in HBS solution of (in mM) 145 NaCl, 22 KCl, 5 glucose, 10 HEPES (pH 7.3) supplemented with 1 CaCl_2_, 0.5 EDTA, 3 NaOH, 0.2 mg/mL cysteine, 1 mg/mL papain and 50 µl/mL Deoxyribonuclease I (Worthington). The tissue was washed with HBS and triturated in plating medium made of MEM with 10% fetal bovine serum, 2 mM GlutaMax. The dissociated cells were pelleted by centrifugation at 1100 rpm for 5 min and suspended in plating medium to count viable cells by trypan blue dye exclusion. For Ca^2+^ imaging, 60,000 to 90,000 cells were plated onto 35 mm dishes with glass bottom (14 mm Dia., No. 0; MatTek Life Sci.) coated with poly-L-lysine (Sigma Aldrich) and incubated for 1 h at 37°C before replacing the plating medium with growth medium made of Neurobasal-A without phenol red supplemented with 2 mM GlutaMax (Gibco) and 2% NeuroCult SM1 Supplement (STEMCELL Tech.). For immunofluorescence, 70,000 cells were plated onto poly-L-lysine-coated glass covers (12 mm Dia., No. 1) in 24-well plates and incubated for 1 h before replacing the medium. At d.i.v. 2, a mix of 37 mM uridine and 27 mM 5-fluoro-2-deoxyuridine (Sigma Aldrich) was added to the cultures.

### Calcium imaging

Membrane-permeant Fura-2 acetoxymethyl (AM) ester (Life Technologies, Eugene OR) suspended in DMSO was added at 4 µM for 1 h at 37°C to neurons in growth medium plated onto 35 mm dishes with manually gridded glass bottom. Cells were washed twice with modified Krebs buffer made of (in mM) 140 NaCl, 5 KCl, 2 CsCl, 5 HEPES, 5 Glucose, 2 Na-Pyruvate, 1 MgCl_2_, 2 CaCl2 (pH 7.4) and incubated with fresh buffer for 15 min at 37°C before transferring to an imaging platform chamber (Warner Instruments, Holliston MA) mounted on an Olympus IX70 inverted microscope equipped with 40x UAPO340 (1.35 N.A.) and 20x UPlanAPO (0.7 N.A.) objectives. The perfusion flow rate (2 ml/min) was controlled with a PPS6 Peristaltic Pump Perfusion System (Warner Instruments). Cells were superfused with modified Krebs buffer containing 0.5 µM tetrodotoxin (TTX; HelloBio) and 1 µM atropine to capture baseline Ca^2+^, followed by co-application for 30 s of 100 nM PNU-282987, 3 µM PNU-120596 (Cayman Chemical), 0.5 µM TTX, 1 µM atropine and wash with 0.5 µM TTX, 1 µM atropine for 12-13 min. Ca^2+^ imaging was accomplished with a Lambda DG-4 illumination system (Sutter Instruments Inc.) enabling rapid switching between 340 and 380 nm excitation wavelengths with emission monitored at 510 nm. Images were acquired with an ORCA digital CMOS camera (Hamamatsu Photonics) controlled with MetaFluor 7.8.13, 64-bit (Molecular Devices, San Jose, CA). For data analysis, F340/F380 ratiometric Ca^2+^ data were exported to Excel and analyzed with OriginPro (OriginLab). For each response trace, the baseline was subtracted by manual correction using two or more points and subtracted values were graphed and analyzed with the Batch Peak Analysis tool (5 cells per batch) to calculate the integrated area under the curve of the traces for the duration of the response and identify maximum peaks. To compare response duration between groups, the integrated area in the first 100 s was calculated and used to derive a Response Decline Index:

*Integrated Area first 100 s/Integrated Area of the entire response*.

For rapidly declining responses, *e.g.* within 100 s, the Response Decline Index is = 1 whereas for protracted responses > 100 s the Index is < 1.

### Immunofluorescence

Cells were fixed with 4% paraformaldehyde (PFA) for 10 min at room temperature (RT) and permeabilized with 0.02% Triton X-100, 3% bovine serum albumin (BSA) in phosphate-buffered saline (PBS) for 10 min at RT and blocked with 3% BSA for 1 h at RT. Primary antibodies in 1% BSA were applied for 1 h at RT followed by three 5 min washes with PBS. Secondary antibodies in 1% BSA were applied for 1 h at RT followed by washes as above. ProLong Gold antifade (Cell Signaling Technology) was added to the glass bottom of dishes with a coverslip placed on top, sandwiching the cells. For immunolabeling of surface Ly6H, fixed cells were incubated with blocking solution followed by incubation for ∼ 16 h at 4°C with anti-Ly6H in 1% BSA. After washing with PBS, cells were permeabilized for 10 min and incubated with blocking solution for 1 h. Primary antibodies (*e.g*. anti-MAP2, GAD65) were diluted in 1% BSA and applied for 1 h at RT followed by incubation with secondary antibodies and mounting with antifade. For tissue labeling, mouse brains were fixed with 4% PFA for ∼16 h at 4°C and washed with PBS 5 times for 10 min before embedding in an agarose/sucrose solution of 3% agarose, 7% sucrose (w/v) in PBS. Coronal slices (30 μm-thick) were cut with a vibratome (Leica Biosystems VT1000S), permeabilized with 0.02% TritonX-100 for 15 min and washed with PBS four times for 5 min. To quench endogenous lipofuscin, the slices were incubated for 45 s with TrueBlack (Biotium) diluted 1:20 in 70% ethanol and immediately washed with PBS. The slices were blocked with Duolink Blocking Solution (Millipore Sigma) for 30 min at RT and incubated with anti-Ly6H and anti-MAP2 for ∼16 h at 4°C. After washing four times with PBS, the slices were incubated for 30 min at RT with donkey anti-mouse secondary antibody conjugated to Cy3 and anti-chicken conjugated to Alexa Fluor 647 in 10% normal donkey serum. After washing four times for 5 min with PBS, the slices were mounted on glass slides with ProLong Gold with DAPI (Cell Signaling Technology).

### α7 nAChR labeling with α-Bungarotoxin

Cells were incubated for 1 h at RT with blocking solution of 3% BSA and for ∼ 16 h at 4°C with α-Bungarotoxin-488 (αBGTX-488; Biotium) diluted 1/500 in 1% BSA in PBS. Cells were washed with PBS three times for 5 min and permeabilized with 0.02% Triton X-100, 3% BSA for 10 min followed by incubation with blocking solution for 1 h. Primary antibodies in 1% BSA were applied for 1 h at RT followed by three 5 min washes with PBS and incubation with secondary antibodies for 1 h. Cells were washed three times for 5 min with PBS and mounted with antifade with DAPI as above.

### Microscopy and analysis of fluorescence

Confocal microscopy was carried out with a Zeiss LSM880 Airyscan using a Plan-Apochromat 63x, 1.4 N.A., oil immersion objective. Images (1024×1024-pixel resolution) were acquired at scan speed set at 8 and pinhole configured to 1 Airy unit for each channel. Image stacks with a 0.5 μm *Z* step were processed and analyzed with Fiji (*76*). For quantitative analysis of Ly6H fluorescence in primary neurons, *Z*-stacks were reconstructed using maximum intensity projection and background subtracted with threshold set at a fixed value of 17. Images were merged with DAPI/MAP2 *Z*-stacks and converted to RGB. MAP2 brightness was increased to better visualize dendrites and soma. A region of interest (R.O.I) around the soma was outlined in the MAP2 image using the segmented line tool and saved. The Analyze Particle function (size = 0.01) was used to measure Ly6H puncta area in the soma R.O.I and normalized to soma area. Primary dendrites connected to the soma were selected with the segmented line tool, straightened (line width 93) and segmented into 10 μm units. Within each segment, the Ly6H images were set to a fixed threshold and converted to mask whereas MAP2 images were automatically thresholded with Li. The processed images were overlayed and Ly6H puncta area measured in each segment with the Analyze Particle tool and normalized to MAP2 area. For quantification of α-Bungarotoxin labeling, αBGTX-488 puncta were manually counted and normalized to the somatodendritic area defined by MAP2 immunolabeling. For tissue slices, images (1912×1912-pixel resolution) were acquired with a Leica SP5 confocal microscope mounted with a 63x oil immersion objective (N.A. = 1.4) at scan speed set at 7 and pinhole configured to 1 Airy unit. Stacks of images were acquired with a 0.6 μm *Z* step. For quantification, maximum projections of Z stack were background subtracted and thresholded with MaxEntropy. An R.O.I of fixed dimensions (20 x 40 µm) was used to sample multiple areas of the CA1 pyramidal layer and *stratum radiatum* for each image. Epifluorescence imaging was carried out using a 20x (N.A. = 1.3) objective or a 60x (N.A. = 1.35) oil immersion objective mounted on a motorized Olympus IX81 inverted microscope equipped with CCD ORCA-R2 camera (Hamamatsu).

### Biotinylation of hippocampal slices

Hippocampi were dissected from P7-8 pups, transferred to cold Hibernate A medium and kept on ice. Tissue sections (400 µm-thick) prepared with a McIlwain Tissue Chopper were transferred to cold External solution of (in mM) 145 NaCl, 2.5 KCl, 2 CaCl_2_ 10 glucose, 10 HEPES (pH 7.4) on ice and separated into floating slices. For each mouse, slices from both hippocampi were washed two times with cold External solution and incubated on ice for 30 min with 1 mg/ml Sulfo-NHS-LC-biotin (Pierce) while slowly rocked. The slices were washed two times with 20 mM Tris in External Solution (pH 7.5), flash-frozen, and stored at –80°C until use. The slices were manually homogenized in a modified RIPA buffer of (in mM) 150 NaCl, 2 EDTA, 50 Tris-HCl (pH 7.4) supplemented with 1% Triton X-100, 0.5% Na deoxycholate, 0.1% SDS, and protease inhibitors (Pierce) and centrifuged at 21,000 x *g* for 20 min at 4°C. Protein content in the supernatant was quantified by Bradford assay (Bio-Rad Labs) and equal amounts of protein per sample mixed with 80 µl of Streptavidin-agarose resin (ThermoScientific) and incubated with rotation for 2 h at 4°C. The resin was washed once with lysis buffer, two times with 0.5% Triton X-100 in phosphate buffer (PB) and two times with PB, with centrifugation at 500 x *g* for 3 min at 4°C between washes. Bound proteins were eluted in SDS-PAGE sample buffer with 100 mM DTT and heated at 95°C for 5 min along with total input protein samples.

### Western blot of tissue and neuron extracts

Mouse hippocampi were homogenized with a mechanical tissue homogenizer in a lysis buffer of (in mM) 150 NaCl, 1 EDTA, 50 Tris-HCl (pH 7.4) supplemented with 1% Triton X-100, 0.5% CHAPS and a cocktail of protease inhibitors. The homogenates were centrifuged at 800 x *g* for 10 min at 4°C and the supernatant collected and centrifuged at 10,000 x *g* for 15 min with the resulting pellet resuspended in lysis buffer. For cell extracts, hippocampal neurons (see Primary cultures) in 24-multiwell plates (200,000 cells/well) were solubilized on ice in modified RIPA buffer and centrifuged at 21,000 x *g* for 20 min at 4°C. For Western blot, proteins were transferred onto nitrocellulose membranes (0.2 µm pore size; Bio-Rad Labs) according to standard procedures. The blocking buffer made in TBST contained 5% BSA for Ly6H and pan-Actin and 5% non-fat dry milk for α7 nAChR and γ-Tubulin. Detection was carried out with horseradish peroxidase (HRP)-conjugated secondary antibodies and chemiluminescence (Immobilon; MilliporeSigma) imaged with Azure c600 (Azure Biosystems).

### Antibodies

Primary antibodies and use conditions are as follows:

|  |  |  |  |
| --- | --- | --- | --- |
| Ly6H/ Mouse | Abnova | AB_425531 | IF 1:400; WB 1:1000 |
| GAD65/ Mouse | DSHB | AB_2314499 | IF 1:400 |
| γ-Tubulin/ Mouse | Sigma Aldrich | AB_477584 | WB 1:2500 |
| Pan-Actin/ Rabbit | Cell Signaling Tech. | AB_10998774 | WB 1:1000 |
| α7 nAChR/ Rabbit | Alomone Labs | AB_10659339 | WB 1:500 |
| GABA/ Rabbit | Sigma Aldrich | AB_477652 | IF 1/8000 |
| MAP2/ Chicken | PhosphoSolutions | AB_2492141 | IF 1:400-500 (slices-cells) |

Secondary antibodies were used according to manufacturer’s specification. For immunofluorescence they included donkey anti-mouse and anti-rabbit conjugated to AlexaFluor-488, –647 (Invitrogen), Cy3 (Jackson ImmunoResearch Labs) and anti-chicken conjugated to AlexaFluor-647, aminomethylcoumarin acetate (Jackson ImmunoResearch Labs). Western blot used goat anti-mouse and anti-rabbit conjugated to HRP (Jackson ImmunoResearch Labs).

### Adenovirus production and transduction

Constructs encoding an H1-pTrip-EF1a-eGFP cassette and Ly6H shRNA or control scrambled shRNA cloned in the pAd/PL-DEST vector were a generous gift from Dr. William J. Joiner (UC San Diego, CA, USA). The constructs were linearized by digestion with Pac I, purified, and transfected with Lipofectamine 3000 (Invitrogen) according to the manufacturer’s protocol in 293A cells plated in 6-well dish at 1.5 x 10^6^ cells/well. Cells were maintained for 10-14 days in I-MEM containing 10% fetal bovine serum, 2 mM glutamine, 10 mM HEPES, 100 U/ml penicillin, 100 µg/ml streptomycin. After collecting the supernatant, cells suspended in PBS were disrupted with three rounds of freeze-thaw and pelleted. One-seventh of the combined supernatants was used to infect 293A cells in 100 mm dishes that were harvested after 4-6 days (1-2 rounds of amplification). The concentration of viral particles was determined by measuring absorbance at 260 nm (OD_260_) to calculate Optical Viral Particle (OVP)/ml: OD_260_ x 0.005 (dilution factor) x 1.1×10^12^ OVP. Viral particles were flash frozen and stored at –80°C until use. For knockdown, neurons from P0/1 pups derived from *Fmr1*^KO^ crossings were virally transduced at d.i.v. 2 with multiplicity of infection (M.O.I.) 20-30:1 based on initial plating density. Viral particles diluted in fresh growth medium (see Primary hippocampal cultures) were added to the cells and incubated for 8 h at 37°C. After infection, the medium was replaced with a mixture of conditioned and fresh growth medium containing uridine/5-fluoro-2-deoxyuridine. Neurons were maintained in culture for 7 days until analysis at d.i.v. 9.

### *In vivo* delivery of PNU-120596

Male and female mice from WT or *Fmr1*^KO^ crossings were used for drug treatment for behavioral studies. Mice were weighed at P17 to determine dose/volume of PNU-120596 for intraperitoneal (i.p.) injection calculated from the average weight of the litter according to the formula V_inj_= ([D] x m_mouse_)/C_1_, where V_inj_ is the injection volume, D is the final concentration of drug in the animal, m_mouse_ is the average weight of the litter in kg, and C_1_ is the working concentration of PNU-120596 injected calculated to ensure that no more than ∼10 μl/g body weight was injected. PNU-120596 (Cayman Chemical) was dissolved in DMSO at 100 mM (31.172 mg/ml) and diluted on the day of the injection to a working concentration of 0.3 mg/ml in filter-sterilized vehicle made of 10% DMSO, 90% corn oil (Sigma Life Sciences). PNU-120596 or vehicle was back filled into a 1 mL syringe (Slip Tip with Intradermal Bevel Needle, 26G x 3/8; BD Syringe) and delivered daily by i.p. injection from P18 to P22, alternating the side of injection each day. Injections were performed blind to genotype and treatment.

### Object Location Memory (OLM) test

Mice were group-housed and randomly assigned to testing sessions; all cages/experimental groups were blinded for both genotype and treatment. Injected mice were habituated to the testing room for 1 h before behavioral assessment. The testing arena was surmounted by a camera for videorecording. OLM was evaluated in an arena with constant visual cues following a standard paradigm consisting of three days of habituation (10 min per trial) followed on day 4 by a training session with objects (10 min) and a test session (5 min) on day 5, when one of the objects was displaced (novel location). Symmetrical rearrangements were avoided to ensure a clear spatial novelty cue and objects were alternatively displaced to rule out intrinsic preference for a given side and or an object. Experiments were performed under controlled acoustic (80 dB) and lighting (1000 lux, 3200K) conditions. Arenas and objects were cleaned with 70% ethanol between subjects. Mouse behavior was analyzed with EthoVision software (Noldus). To assess object location memory, we used a Novel Exploratory Preference Score (EPS) defined as the proportion of time each animal spent exploring the novel object relative to its total object exploration time. To account for potential sex differences (*46, 77*), Z-scores were calculated for each individual based on the mean and standard deviation of the WT control group, stratified by sex. The Z-score normalization used the formula:

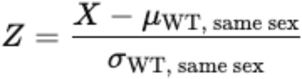

where X is the EPS of an individual animal, and μ and σ represent the mean and standard deviation of the corresponding sex-specific WT control group. Animals with a Z-score ≥ –1 were considered to have passed the task (Pass), reflecting behavior consistent with intact object location memory, *e.g.* a measurable preference for the novel location. Conversely, animals with Z-scores < –1 were classified as failing to demonstrate this preference (Fail), indicating impaired spatial memory or reduced novelty discrimination. This threshold represents a conservative criterion, identifying individuals that perform more than one standard deviation below the WT average as behaviorally impaired, and enables a standardized comparison across experimental groups.

### Hyperactivity

Drug-treated mice at age P22 were transferred from the animal facility and allowed to habituate to the laboratory room for > 30 min. All litters were tested in a dedicated, isolated room starting in mid-afternoon, to minimize possible circadian effects. The testing chamber was surmounted by a camera for videorecording. Mice were tracked for a total of 10 min, 5 min of habituation followed by 5 min of activity measurement. Mouse behavior was analyzed with EthoVision software.

### Audiogenic seizure (AGS)

AGS was induced with an alarm (Portable Safety Kit Door Alarm; RadioShack) set at ∼120 dB. The alarm was turned on for 60 s and the seizure behavior videorecorded. AGS was analyzed blind to drug treatment utilizing the video recording. AGS was scored using the Audiogenic Response Score that is specifically designed to capture the signs and severity of these generalized motor seizures (*55*).

### Statistical analysis

Data were analyzed by GraphPad Prism 8.1 and JMP Pro 18.0.2 software. Unless otherwise indicated, statistical significance was determined by t-test (two-tailed) or one-way analysis of variance (ANOVA) (Tukey’s multiple-comparison post-test), with p < 0.05 considered significant.

## SUPPLEMENTAL MATERIAL

**Figure S1**. Ly6H protein is robustly expressed in WT hippocampus during the first week of postnatal development.

**Figure S2**. Ly6H protein expression is not altered in primary *Fmr1*^KO^ hippocampal neurons.

**Figure S3.** α7 nAChR agonist sensitivity in WT primary hippocampal neurons.

**Figure S4.** α7 nAChR expression is not altered in immature *Fmr1*^KO^ hippocampal neurons.

**Figure S5.** Reduced expression of Ly6H in hippocampal neurons transduced with Ly6H shRNA.

**Figure S6.** Impaired spatial memory in naïve early adolescent *Fmr1*^KO^ mice.

**Figure S7.** Hyperlocomotor activity in naïve early adolescent *Fmr1*^KO^ mice.

**Figure S8.** AGS progression and scoring system.

## Acknowledgments

We thank the members of the Francesconi laboratory for assistance with methodology and critical insights. We thank Noelie Cayla for assistance with EthoVision. We acknowledge the assistance of the Neural Cell Engineering and Imaging Core of the Einstein Rose F. Kennedy Intellectual and Developmental Disabilities Research Center supported by the National Institute of Child Health and Human Development U54 HD090260.

## Funding

National Institutes of Health grant U54 HD090260 (Walkley SU, PI); IDDRC Pilot Project (AF)

National Institutes of Health grant MH108614 (AF)

National Institutes of Health grant S10OD025295 (Instrumentation grant; Dobrenis K, PI)

## Author contributions

Conceptualization: SG, VKV, AF

Methodology: SG, DCM, VKV, AF

Investigation: SG, DCM, AF

Visualization: SG, VKV, AF

Funding acquisition: VKV, AF

Project administration: AF

Supervision: VKV, AF

Writing – original draft: SG

Writing – review & editing: SG, VKV, AF

## Competing interests

Authors declare that they have no competing interests.

## Data and materials availability

All data generated or analyzed during this study are included in the manuscript and supporting files.

## Supplementary Material for

**The Supplementary Material includes**:

Figures S1 to S8.

**Figure S1.**
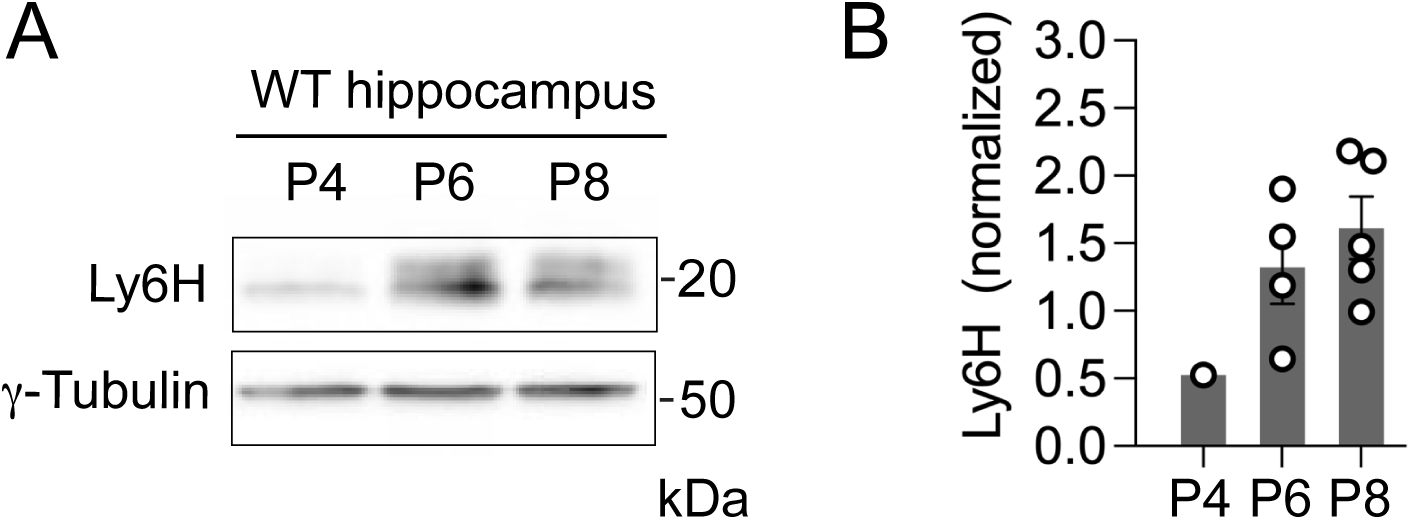
Ly6H protein is robustly expressed in WT hippocampus during the first week of postnatal development. **(A)** Representative Western blot of WT hippocampal tissue lysates at P4, P6, and P8 probed for Ly6H and γ-Tubulin (loading control). **(B)** Ly6H expression relative to γ-Tubulin: mean ± SEM, P4 0.526 n = 1, P6 1.32 ± 0.268 n = 4, P8 1.61 ± 0.232 n = 5 mice.

**Figure S2.**
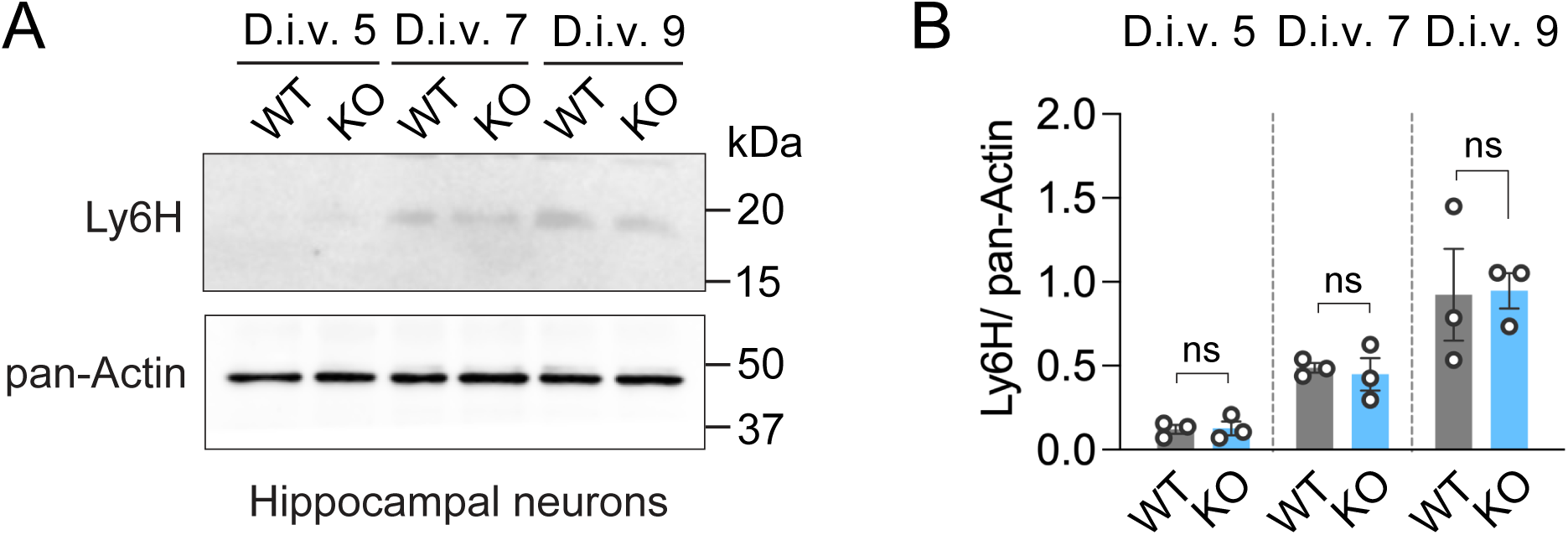
Ly6H protein expression is not altered in primary *Fmr1*^KO^ hippocampal neurons. **(A**) Western blot of extracts from WT and *Fmr1*^KO^ hippocampal neurons at different ages *in vitro* probed for Ly6H and pan-Actin (loading control). (**B**) Ly6H relative to Actin. Mean ± SEM WT d.i.v. 5, 0.121 ± 0.0265; d.i.v. 7, 0.487 ± 0.0291; d.i.v. 9, 0.924 ± 0.273; KO d.i.v. 5, 0.127 ± 0.0414; d.i.v. 7, 0.449 ± 0.0962; d.i.v. 9, 0.947 ± 0.106, N = 3 independent cultures, unpaired t-test.

**Figure S3.**
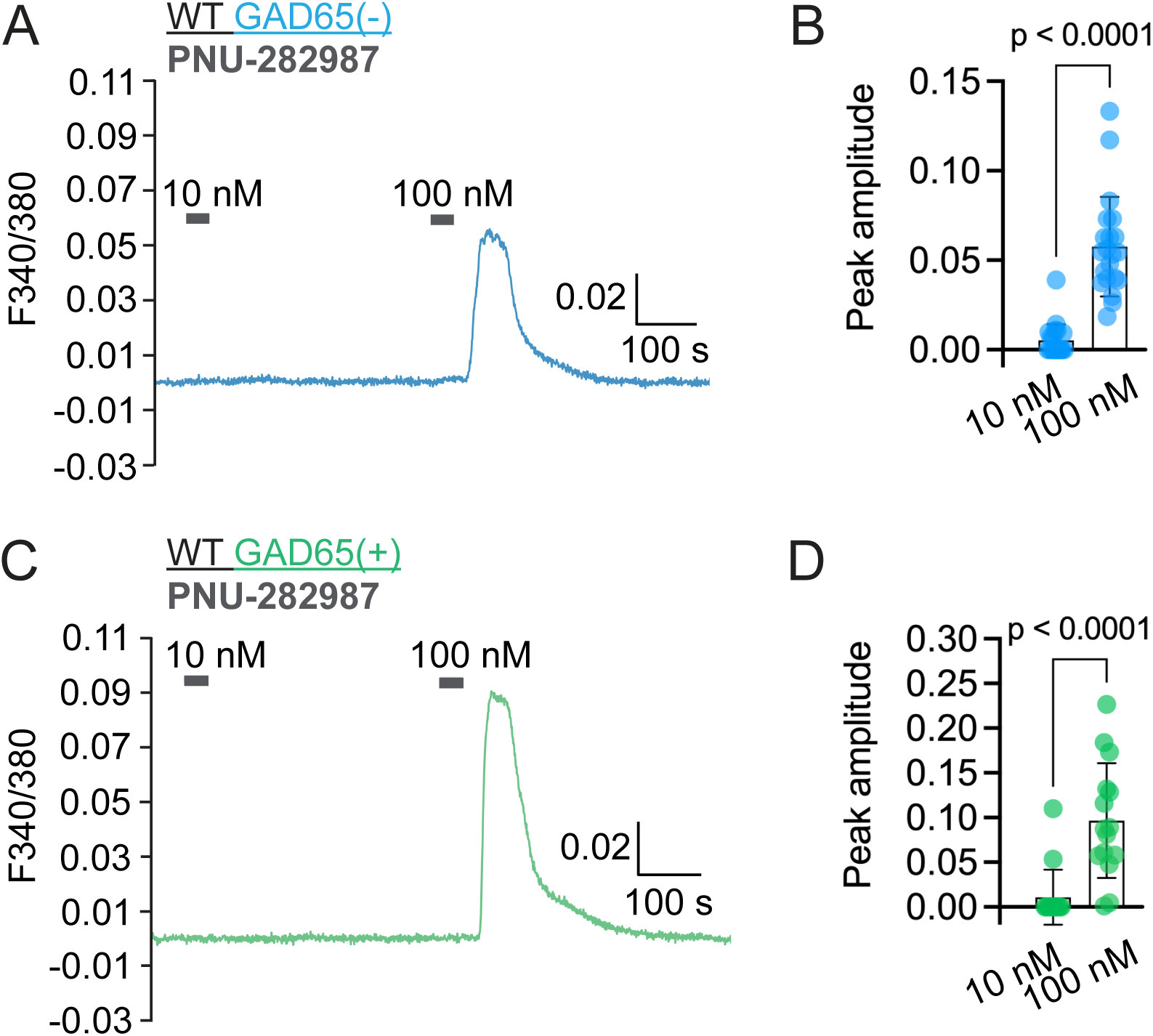
α7 nAChR agonist sensitivity in WT primary hippocampal neurons. (**A**) Representative traces of Ca^2+^ responses evoked by 10 nM and 100 nM PNU-282987 with 3 µM PNU-120596 (gray bars) in WT d.i.v. 7 GAD65(−) neurons. (**B**) Quantification of peak amplitude of evoked responses in GAD65(−) neurons. Mean ± SEM, 10 nM 0.00511 ± 0.002, 100 nM 0.0576 ± 0.00607, n = 21 per group, Mann-Whitney test. (**C**) Representative traces of Ca^2+^ responses evoked as above in WT d.i.v. 7 GAD65(+) neurons. (**D**) Quantification of peak amplitude in GAD65(+) cells. Mean ± SEM, 10 nM 0.0109 ± 0.0965, 100 nM 0.0965 ± 0.0166, n = 15 per group, Mann-Whitney test.

**Figure S4.**
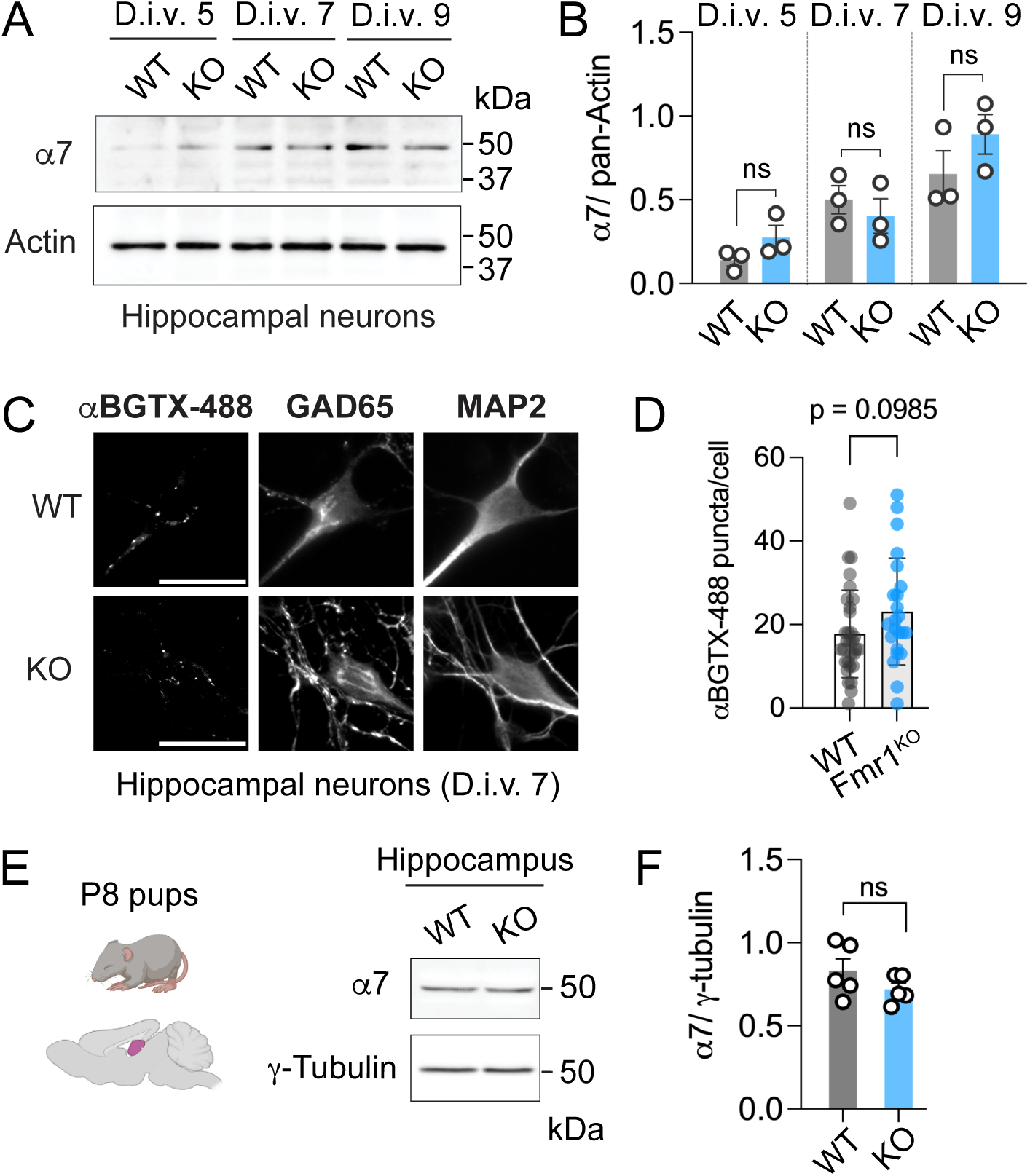
α7 nAChR expression is not altered in immature *Fmr1*^KO^ hippocampal neurons. **(A)** Representative Western blot of primary hippocampal neurons at different ages *in vitro* probed for α7 and pan-Actin (loading control as in Figure S2). (**B**) Quantification of α7 subunit relative to Actin. Mean ± SEM WT d.i.v. 5, 0.14 ± 0.035; d.i.v. 7, 0.5 ± 0.084; d.i.v. 9, 0.654 ± 0.139; KO d.i.v. 5, 0.275 ± 0.0717; d.i.v. 7, 0.402 ± 0.103; d.i.v. 9, 0.891 ± 0.118, N = 3 independent cultures, unpaired t-test. (**C**) Representative images of bound αBGTX-488 in non-permeabilized neurons co-labeled for GAD65 and MAP2; scale bar, 25 µm. (**D**) Quantification of αBGTX-488 clusters in the somatodendritic region of GAD65(+) neurons. Mean ± SEM, WT 17.7 ± 1.89, n = 31, KO 23.1 ± 2.67 n = 23 neurons from N = 3 independent experiments, unpaired t-test. (**E**) Representative Western blot of extracts from WT and *Fmr1*^KO^ P8 hippocampus probed for α7 and ψ-Tubulin (loading control). (**F**) Quantification of α7 abundance relative to ψ-Tubulin. Mean ± SEM, WT 0.832 ± 0.0718, KO 0.719 ± 0.0369, n = 5 mice per group, unpaired t-test.

**Figure S5.**
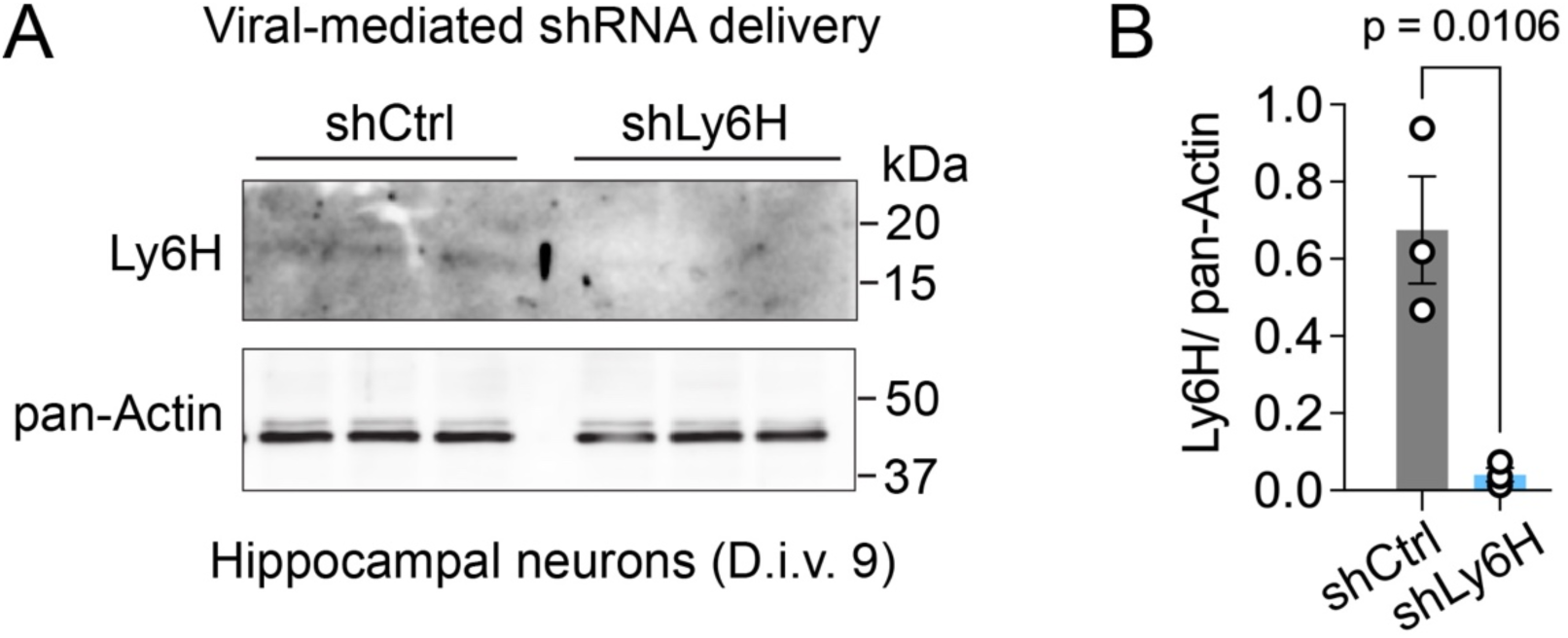
Reduced expression of Ly6H in hippocampal neurons transduced with Ly6H shRNA. **(A**) Representative Western blot of d.i.v. 9 hippocampal neurons transduced with control (shCtrl) or Ly6H shRNA (shLy6H) probed for Ly6H and pan-Actin (loading control). (**B**) Quantification of Ly6H relative to Actin. Mean ± SEM, control 0.68 ± 0.14, Ly6H shRNA 0.041 ± 0.018, N = 3, unpaired t-test.

**Figure S6.**
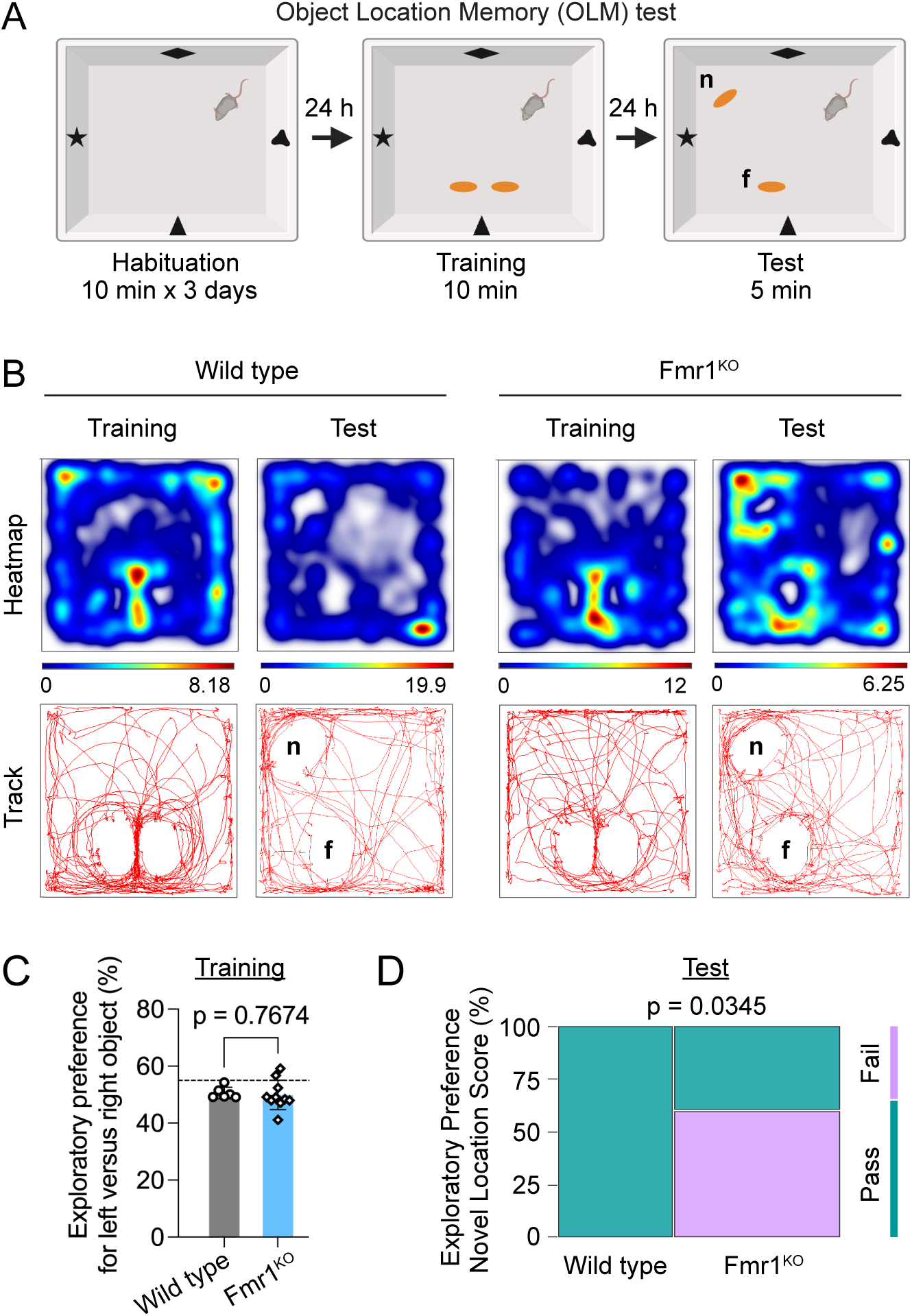
Impaired spatial memory in naïve early adolescent *Fmr1*^KO^ mice. **(A)** Schematic and timeline of OLM test: *n* indicates the novel location of the displaced object, and *f* the familiar location of the stationary one (created with BioRender). (**B**) Representative activity heatmaps and tracks for P22 WT and *Fmr1*^KO^ mice during training (last 5 min) and test phase (5 min). (**C**) Exploratory preference for individual objects (left *vs*. right) during the training phase with 50% indicating no preference. Mean ± SEM, WT 50.6 ± 0.82% n = 6, *Fmr1*^KO^ 49.9 ± 1.62% n = 10, unpaired t-test. (**D**) Mosaic plot of Exploratory Preference Score expressed as Z-score stratified by sex, where preference for novel location (Pass) corresponds to Z ≥ –1; WT n = 7, *Fmr1*^KO^ n = 10 mice; Fisher’s Exact Test.

**Figure S7.**
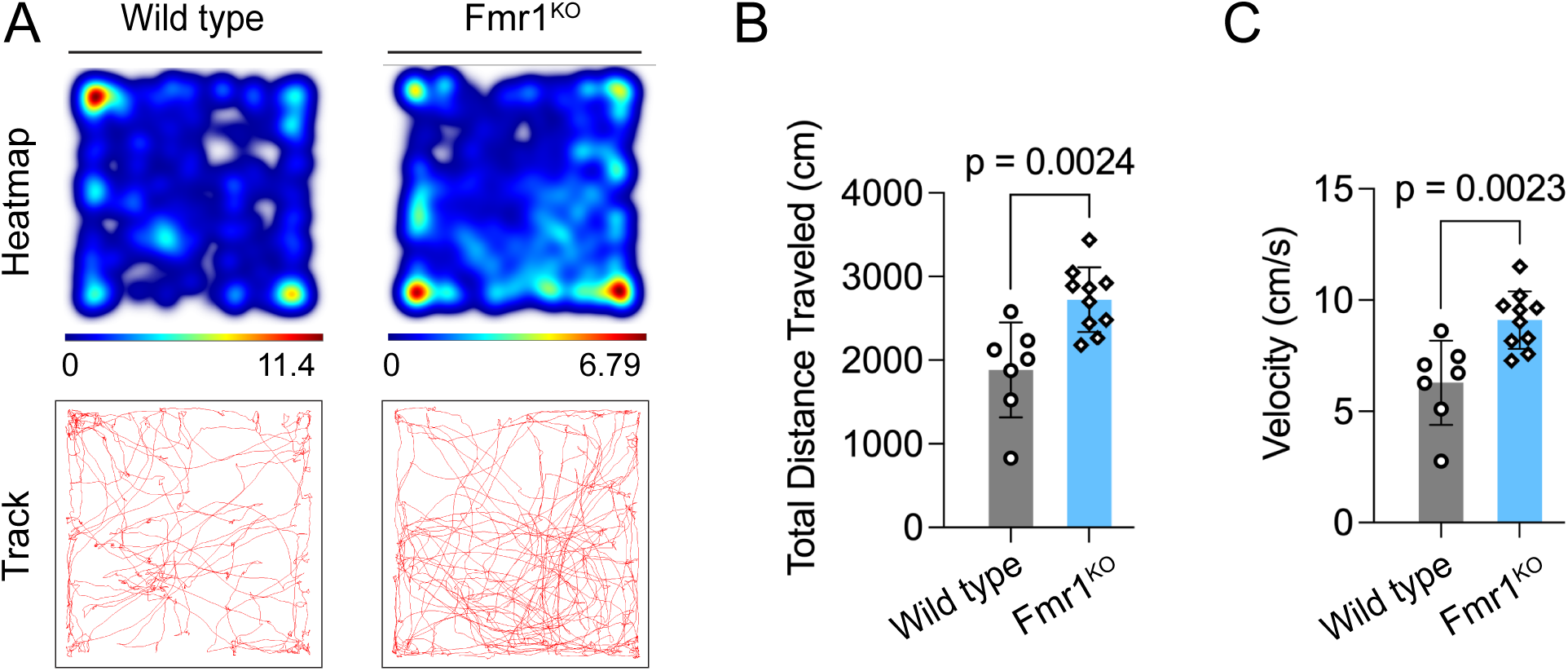
Hyperlocomotor activity in naïve early adolescent *Fmr1*^KO^ mice. **(A)** Representative activity heatmaps and tracks in open field of P22 WT and *Fmr1*^KO^ mice after one day of habituation. **(B)** Total distance traveled: mean ± SEM, WT vehicle 1883 ± 214 cm n = 7, *Fmr1*^KO^ 2723 ± 122 n = 10 mice, unpaired t-test. (**C**) Velocity: mean ± SEM, WT vehicle 6.29 ± 0.715 cm/s n = 7, *Fmr1*^KO^ 9.1 ± 0.41 n = 10 mice, unpaired t-test.

**Figure S8.**
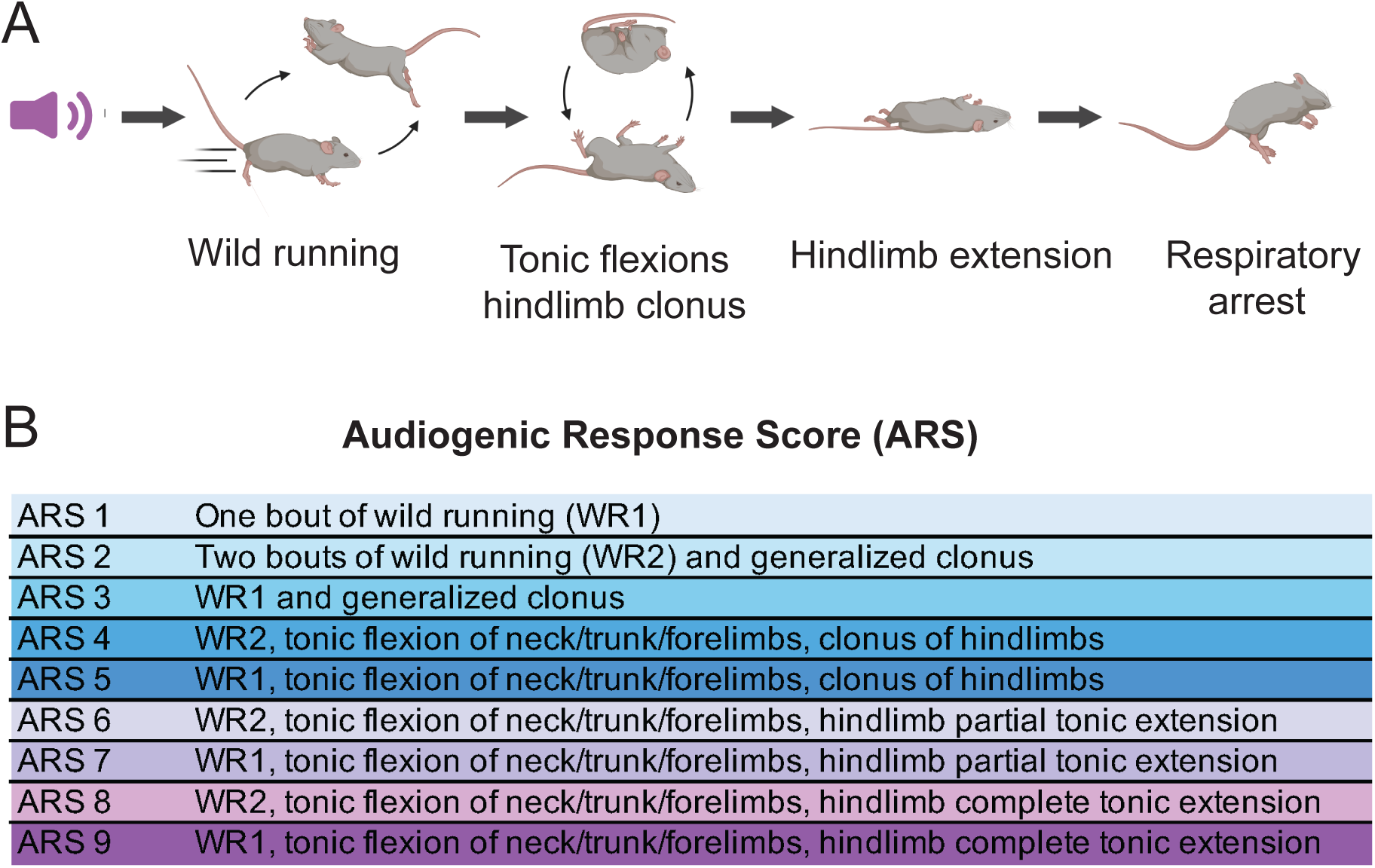
AGS progression and scoring system. (**A**) Illustration of the progression of audiogenic motor seizures (created with BioRender). (**B**) Description of Audiogenic Response Score (ARS) criteria used in the study.

## Notes

### Competing Interest Statement

The authors have declared no competing interest.

### Summary of Updates

New results as reported in Fig 6, Fig S1B, Fig S3, Fig S6, and Fig S7. Updated Fig S4 now includes data from original Fig S3. Manuscript title, abstract and main text were revised to include new findings.

## REFERENCES

1. J. Hunter et al., Epidemiology of fragile X syndrome: a systematic review and meta-analysis. Am J Med Genet A 164a, 1648–1658 (2014).

2. D. B. Bailey, Jr., M. Raspa, M. Olmsted, D. B. Holiday, Co-occurring conditions associated with FMR1 gene variations: findings from a national parent survey. Am J Med Genet A 146A, 2060–2069 (2008).

3. W. E. Kaufmann et al., Autism Spectrum Disorder in Fragile X Syndrome: Cooccurring Conditions and Current Treatment. Pediatrics 139, S194–S206 (2017).

4. A. J. Verkerk et al., Identification of a gene (FMR-1) containing a CGG repeat coincident with a breakpoint cluster region exhibiting length variation in fragile X syndrome. Cell 65, 905–914 (1991).

5. J. D. Richter, X. Zhao, The molecular biology of FMRP: new insights into fragile X syndrome. Nat Rev Neurosci 22, 209–222 (2021).

6. X. Liu, V. Kumar, N. P. Tsai, B. D. Auerbach, Hyperexcitability and Homeostasis in Fragile X Syndrome. Front Mol Neurosci 14, 805929 (2021).

7. L. Maggi, E. Sher, E. Cherubini, Regulation of GABA release by nicotinic acetylcholine receptors in the neonatal rat hippocampus. J Physiol 536, 89–100 (2001).

8. L. M. Teles-Grilo Ruivo, J. R. Mellor, Cholinergic modulation of hippocampal network function. Front Synaptic Neurosci 5, 2 (2013).

9. J. L. Yakel, Nicotinic ACh receptors in the hippocampal circuit; functional expression and role in synaptic plasticity. J Physiol 592, 4147–4153 (2014).

10. L. Maggi, C. Le Magueresse, J. P. Changeux, E. Cherubini, Nicotine activates immature “silent” connections in the developing hippocampus. Proc Natl Acad Sci U S A 100, 2059–2064 (2003).

11. Z. Liu, R. A. Neff, D. K. Berg, Sequential interplay of nicotinic and GABAergic signaling guides neuronal development. Science 314, 1610–1613 (2006).

12. A. F. Lozada et al., Glutamatergic synapse formation is promoted by alpha7-containing nicotinic acetylcholine receptors. J Neurosci 32, 7651–7661 (2012).

13. H. Lin et al., Cortical parvalbumin GABAergic deficits with alpha7 nicotinic acetylcholine receptor deletion: implications for schizophrenia. Mol Cell Neurosci 61, 163–175 (2014).

14. L. J. Volk, B. E. Pfeiffer, J. R. Gibson, K. M. Huber, Multiple Gq-coupled receptors converge on a common protein synthesis-dependent long-term depression that is affected in fragile X syndrome mental retardation. J Neurosci 27, 11624–11634 (2007).

15. S. R. Thomson et al., Cell-Type-Specific Translation Profiling Reveals a Novel Strategy for Treating Fragile X Syndrome. Neuron 95, 550–563 e555 (2017).

16. R. S. Hurst et al., A novel positive allosteric modulator of the alpha7 neuronal nicotinic acetylcholine receptor: in vitro and in vivo characterization. J Neurosci 25, 4396–4405 (2005).

17. J. W. Young et al., Impaired attention is central to the cognitive deficits observed in alpha 7 deficient mice. Eur Neuropsychopharmacol 17, 145–155 (2007).

18. A. J. Sharp et al., A recurrent 15q13.3 microdeletion syndrome associated with mental retardation and seizures. Nat Genet 40, 322–328 (2008).

19. C. E. Adams et al., Reduced Chrna7 expression in mice is associated with decreases in hippocampal markers of inhibitory function: implications for neuropsychiatric diseases. Neuroscience 207, 274–282 (2012).

20. S. Fucile, Ca2+ permeability of nicotinic acetylcholine receptors. Cell Calcium 35, 1–8 (2004).

21. S. F. Colombo, F. Mazzo, F. Pistillo, C. Gotti, Biogenesis, trafficking and up-regulation of nicotinic ACh receptors. Biochem Pharmacol 86, 1063–1073 (2013).

22. J. A. Matta, S. Gu, W. B. Davini, D. S. Bredt, Nicotinic acetylcholine receptor redux: Discovery of accessories opens therapeutic vistas. Science 373, (2021).

23. J. M. Miwa, K. R. Anderson, K. M. Hoffman, Lynx Prototoxins: Roles of Endogenous Mammalian Neurotoxin-Like Proteins in Modulating Nicotinic Acetylcholine Receptor Function to Influence Complex Biological Processes. Front Pharmacol 10, 343 (2019).

24. J. M. Miwa et al., lynx1, an endogenous toxin-like modulator of nicotinic acetylcholine receptors in the mammalian CNS. Neuron 23, 105–114 (1999).

25. I. Ibanez-Tallon et al., Novel modulation of neuronal nicotinic acetylcholine receptors by association with the endogenous prototoxin lynx1. Neuron 33, 893–903 (2002).

26. C. A. Puddifoot, M. Wu, R. J. Sung, W. J. Joiner, Ly6h regulates trafficking of alpha7 nicotinic acetylcholine receptors and nicotine-induced potentiation of glutamatergic signaling. J Neurosci 35, 3420–3430 (2015).

27. M. Wu, C. A. Puddifoot, P. Taylor, W. J. Joiner, Mechanisms of inhibition and potentiation of alpha4beta2 nicotinic acetylcholine receptors by members of the Ly6 protein family. J Biol Chem 290, 24509–24518 (2015).

28. M. Kalinowska, C. Castillo, A. Francesconi, Quantitative profiling of brain lipid raft proteome in a mouse model of fragile X syndrome. PLoS One 10, e0121464 (2015).

29. Y. Moriwaki et al., Endogenous neurotoxin-like protein Ly6H inhibits alpha7 nicotinic acetylcholine receptor currents at the plasma membrane. Sci Rep 10, 11996 (2020).

30. M. Wu, C. Z. Liu, E. A. Barrall, R. A. Rissman, W. J. Joiner, Unbalanced Regulation of alpha7 nAChRs by Ly6h and NACHO Contributes to Neurotoxicity in Alzheimer’s Disease. J Neurosci 41, 8461–8474 (2021).

31. A. Zeisel et al., Molecular Architecture of the Mouse Nervous System. Cell 174, 999–1014.e1022 (2018).

32. M. S. Cembrowski, L. Wang, K. Sugino, B. C. Shields, N. Spruston, Hipposeq: a comprehensive RNA-seq database of gene expression in hippocampal principal neurons. eLife 5, e14997 (2016).

33. M. Alkondon, E. X. Albuquerque, Diversity of nicotinic acetylcholine receptors in rat hippocampal neurons. I. Pharmacological and functional evidence for distinct structural subtypes. J Pharmacol Exp Ther 265, 1455–1473 (1993).

34. M. S. Thomsen et al., Expression of the Ly-6 family proteins Lynx1 and Ly6H in the rat brain is compartmentalized, cell-type specific, and developmentally regulated. Brain Struct Funct 219, 1923–1934 (2014).

35. R. Freedman, C. Wetmore, I. Stromberg, S. Leonard, L. Olson, Alpha-bungarotoxin binding to hippocampal interneurons: immunocytochemical characterization and effects on growth factor expression. J Neurosci 13, 1965–1975 (1993).

36. C. E. Adams et al., Development of the alpha7 nicotinic cholinergic receptor in rat hippocampal formation. Brain Res Dev Brain Res 139, 175–187 (2002).

37. M. Hajos et al., The selective alpha7 nicotinic acetylcholine receptor agonist PNU-282987 [N-[(3R)-1-Azabicyclo[2.2.2]oct-3-yl]-4-chlorobenzamide hydrochloride] enhances GABAergic synaptic activity in brain slices and restores auditory gating deficits in anesthetized rats. J Pharmacol Exp Ther 312, 1213–1222 (2005).

38. B. I. Kalappa, A. G. Gusev, V. V. Uteshev, Activation of functional α7-containing nAChRs in hippocampal CA1 pyramidal neurons by physiological levels of choline in the presence of PNU-120596. PLoS One 5, e13964 (2010).

39. A. L. Bodnar et al., Discovery and structure-activity relationship of quinuclidine benzamides as agonists of alpha7 nicotinic acetylcholine receptors. J Med Chem 48, 905–908 (2005).

40. K. Ganguly, A. F. Schinder, S. T. Wong, M. Poo, GABA itself promotes the developmental switch of neuronal GABAergic responses from excitation to inhibition. Cell 105, 521–532 (2001).

41. R. Tyzio, G. L. Holmes, Y. Ben-Ari, R. Khazipov, Timing of the developmental switch in GABA(A) mediated signaling from excitation to inhibition in CA3 rat hippocampus using gramicidin perforated patch and extracellular recordings. Epilepsia 48 **Suppl 5**, 96–105 (2007).

42. C. Bagni, R. S. Zukin, A Synaptic Perspective of Fragile X Syndrome and Autism Spectrum Disorders. Neuron 101, 1070–1088 (2019).

43. T. Murai, S. Okuda, T. Tanaka, H. Ohta, Characteristics of object location memory in mice: Behavioral and pharmacological studies. Physiol Behav 90, 116–124 (2007).

44. R. R. Seese, K. Wang, Y. Q. Yao, G. Lynch, C. M. Gall, Spaced training rescues memory and ERK1/2 signaling in fragile X syndrome model mice. Proc Natl Acad Sci U S A 111, 16907–16912 (2014).

45. G. Ntoulas et al., Multi-level profiling of the Fmr1 KO rat unveils altered behavioral traits along with aberrant glutamatergic function. Transl Psychiatry 14, 104 (2024).

46. A. Cruz-Sanchez et al., Developmental onset distinguishes three types of spontaneous recognition memory in mice. Scientific Reports 10, 10612 (2020).

47. Z. Gu, K. D. Stevanovic, J. D. Cushman, J. L. Yakel, Cholinergic-Sensitive Theta Oscillations in Memory Encoding in Mice. The Journal of Neuroscience 44, e1313232024 (2024).

48. Q. J. Yan, M. Rammal, M. Tranfaglia, R. P. Bauchwitz, Suppression of two major Fragile X Syndrome mouse model phenotypes by the mGluR5 antagonist MPEP. Neuropharmacology 49, 1053–1066 (2005).

49. T. M. Kazdoba, P. T. Leach, J. L. Silverman, J. N. Crawley, Modeling fragile X syndrome in the Fmr1 knockout mouse. Intractable Rare Dis Res 3, 118–133 (2014).

50. Y. Zhang et al., Loss of MeCP2 in cholinergic neurons causes part of RTT-like phenotypes via alpha7 receptor in hippocampus. Cell Res 26, 728–742 (2016).

51. N. K. Sharma, S. Kaur, R. K. Goel, Exploring the ameliorative role of alpha7 neuronal nicotinic acetylcholine receptor modulation in epilepsy and associated comorbidities in post-PTZ-kindled mice. Epilepsy Behav 103, 106862 (2020).

52. P. Sun, D. G. Liu, X. M. Ye, Nicotinic Acetylcholine Receptor alpha7 Subunit Is an Essential Regulator of Seizure Susceptibility. Front Neurol 12, 656752 (2021).

53. J. J. Zheng et al., Cytisine Exerts an Anti-Epileptic Effect via alpha7nAChRs in a Rat Model of Temporal Lobe Epilepsy. Front Pharmacol 12, 706225 (2021).

54. M. A. Gillentine, C. P. Schaaf, The human clinical phenotypes of altered CHRNA7 copy number. Biochem Pharmacol 97, 352–362 (2015).

55. P. C. Jobe, A. L. Picchioni, L. Chin, Role of brain 5-hydroxytryptamine in audiogenic seizure in the rat. Life Sci 13, 1–13 (1973).

56. L. G. Romanova, Z. A. Zorina, L. I. Korochkin, A genetic, physiological, and biochemical investigation of audiogenic seizures in rats. Behav Genet 23, 483–489 (1993).

57. K. C. Ross, J. R. Coleman, Developmental and genetic audiogenic seizure models: behavior and biological substrates. Neurosci Biobehav Rev 24, 639–653 (2000).

58. J. C. Darnell et al., FMRP stalls ribosomal translocation on mRNAs linked to synaptic function and autism. Cell 146, 247–261 (2011).

59. T. Maurin et al., HITS-CLIP in various brain areas reveals new targets and new modalities of RNA binding by fragile X mental retardation protein. Nucleic Acids Res 46, 6344–6355 (2018).

60. A. B. Tekinay et al., A role for LYNX2 in anxiety-related behavior. Proc Natl Acad Sci U S A 106, 4477–4482 (2009).

61. E. Dessaud, D. Salaun, O. Gayet, M. Chabbert, O. deLapeyriere, Identification of lynx2, a novel member of the ly-6/neurotoxin superfamily, expressed in neuronal subpopulations during mouse development. Mol Cell Neurosci 31, 232–242 (2006).

62. M. J. Boland et al., Molecular analyses of neurogenic defects in a human pluripotent stem cell model of fragile X syndrome. Brain 140, 582–598 (2017).

63. K. Talvio et al., Reduced LYNX1 expression in transcriptome of human iPSC-derived neural progenitors modeling fragile X syndrome. Front Cell Dev Biol 10, 1034679 (2022).

64. N. R. Campbell, C. C. Fernandes, A. W. Halff, D. K. Berg, Endogenous signaling through alpha7-containing nicotinic receptors promotes maturation and integration of adult-born neurons in the hippocampus. J Neurosci 30, 8734–8744 (2010).

65. Z. Gu et al., Hippocampal Interneuronal alpha7 nAChRs Modulate Theta Oscillations in Freely Moving Mice. Cell Rep 31, 107740 (2020).

66. A. Tempio, A. Boulksibat, B. Bardoni, S. Delhaye, Fragile X Syndrome as an interneuronopathy: a lesson for future studies and treatments. Front Neurosci 17, 1171895 (2023).

67. T. J. Wills, F. Cacucci, N. Burgess, J. O’Keefe, Development of the hippocampal cognitive map in preweanling rats. Science 328, 1573–1576 (2010).

68. N. Jeong, A. C. Singer, Learning from inhibition: Functional roles of hippocampal CA1 inhibition in spatial learning and memory. Curr Opin Neurobiol 76, 102604 (2022).

69. R. J. Hagerman et al., Fragile X syndrome. Nat Rev Dis Primers 3, 17065 (2017).

70. A. P. F. Domanski, S. A. Booker, D. J. A. Wyllie, J. T. R. Isaac, P. C. Kind, Cellular and synaptic phenotypes lead to disrupted information processing in Fmr1-KO mouse layer 4 barrel cortex. Nat Commun 10, 4814 (2019).

71. E. N. Falk et al., Nicotinic regulation of local and long-range input balance drives top-down attentional circuit maturation. Sci Adv 7, (2021).

72. A. Contractor, V. A. Klyachko, C. Portera-Cailliau, Altered Neuronal and Circuit Excitability in Fragile X Syndrome. Neuron 87, 699–715 (2015).

73. C. A. Trujillo et al., Pharmacological reversal of synaptic and network pathology in human MECP2-KO neurons and cortical organoids. EMBO Mol Med 13, e12523 (2021).

74. A. S. Lewis et al., An Exploratory Trial of Transdermal Nicotine for Aggression and Irritability in Adults with Autism Spectrum Disorder. J Autism Dev Disord 48, 2748–2757 (2018).

75. K. W. Gee et al., First in human trial of a type I positive allosteric modulator of alpha7-nicotinic acetylcholine receptors: Pharmacokinetics, safety, and evidence for neurocognitive effect of AVL-3288. J Psychopharmacol 31, 434–441 (2017).

76. J. Schindelin et al., Fiji: an open-source platform for biological-image analysis. Nat Methods 9, 676-682 (2012).

77. L. Ricceri, C. Colozza, G. Calamandrei, Ontogeny of spatial discrimination in mice: a longitudinal analysis in the modified open-field with objects. Dev Psychobiol 37, 109–118 (2000).

